# Synovial Sarcoma Chromatin Dynamics Reveal a Continuum in SS18:SSX Reprograming

**DOI:** 10.1101/2024.05.14.594262

**Authors:** Jakob Hofvander, Alvin Qiu, Kiera Lee, Misha Bilenky, Annaïck Carles, Qi Cao, Michelle Moksa, Jonathan Steif, Edmund Su, Afroditi Sotiriou, Angela Goytain, Lesley A. Hill, Sam Singer, Irene L. Andrulis, Jay S. Wunder, Fredrik Mertens, Ana Banito, Kevin B. Jones, T. Michael Underhill, Torsten O. Nielsen, Martin Hirst

## Abstract

Synovial sarcoma (SyS) is an aggressive soft-tissue malignancy characterized by a pathognomonic chromosomal translocation leading to the formation of the SS18::SSX fusion oncoprotein. SS18::SSX associates with mammalian BAF complexes suggesting deregulation of chromatin architecture as the oncogenic driver in this tumour type. To examine the epigenomic state of SyS we performed comprehensive multi-omics analysis on 52 primary pre-treatment human SyS tumours. Our analysis revealed a continuum of epigenomic states across the cohort at fusion target genes independent of rare somatic genetic lesions. We identify cell-of-origin signatures defined by enhancer states and reveal unexpected relationships between H2AK119Ub1 and active marks. The number of bivalent promoters, dually marked by the repressive H3K27me3 and activating H3K4me3 marks, has strong prognostic value and outperforms tumor grade in predicting patient outcome. Finally, we identify SyS defining epigenomic features including H3K4me3 expansion associated with striking promoter DNA hypomethylation in which SyS displays the lowest mean methylation level of any sarcoma subtype. We explore these distinctive features as potential vulnerabilities in SyS and identify H3K4me3 inhibition as a promising therapeutic strategy.

---

Many types of sarcoma, particularly those affecting younger patients, are considered predominantly epigenetic diseases where widespread epigenetic dysregulation is initiated by a small number of, or even a single, genetic change^1,2^. Among these tumor types are several fusion driven sarcomas^3^, including SyS, one of the most common soft tissue sarcomas in adolescents and young adults^4^. SyS is an aggressive malignancy with poor prognosis^5,6^, few effective treatment options, and a survival rate that has failed to improve over recent decades^7^. SyS is defined by the pathognomonic t(X;18) chromosomal translocation resulting in the formation of the SS18::SSX fusion oncoprotein^8,9^.

SS18 is a known member of the BRM/BRG1 associated factors (BAF) complex, a set of ATP-dependent chromatin remodeling complexes^10,11^. BAF complexes function to alter the nucleosome architecture, creating DNA accessibility at enhancers and promoters to allow for binding of both transcription factors and transcriptional machinery^12,13^. SS18::SSX competes with endogenous SS18 to incorporate into the canonical (cBAF) and non-canonical (GBAF) forms of the remodeling complex^14,15^. Incorporation of the fusion protein into cBAF results in its degradation, which in turn increases the relative prevalence of other BAF-family complexes (polybromo-associated (PBAF) and GBAF), altering the BAF subtype balance and genomic distribution^14^. The observed synthetic lethality associated with elevated prevalence of GBAF indicated a potential therapeutic vulnerability in SyS^16,17^, although recent clinical trials focused on targeting BRD9 (a specific component of the GBAF complex) did not show clinical benefit (NCT05355753; NCT04965753).

SS18::SSX interacts genetically and physically with numerous epigenetic regulatory proteins including the DNA binding protein ATF2, the transcriptional corepressor TLE1 and members of non-canonical polycomb group repressor complexes^18,19^ (PRC1.1, 1.3 and 1.5). SS18::SSX containing BAF complexes display increased nucleosome-binding properties, specifically at H2AK119Ub1-modified nucleosomes, conferred by a direct interaction between SSX and the nucleosome acidic patch^20–22^. H2AK119Ub1 is essential for maintaining transcriptional repression and H3K27me3 deposition^23^; however, recruitment of BAF to these sites results in H3K27me3 eviction and inappropriate gene activation through a mechanism that is not fully understood^20,21,24,25^ . Thus, while multiple mechanisms of action for SS18::SSX have been proposed, they converge on the formation of unusually broad BAF domains that are associated with the loss of H3K27me3 and gain of active histone marks ^18,19,14^.

SyS is genomically stable and displays few somatic mutations other than the chromosomal translocation itself, suggesting that SS18::SSX bears the prime responsibility for malignant transformation^26–28^. Despite this shared molecular basis, the clinical and histological presentation of SyS varies significantly. We thus sought to test if the clinical variability observed in SyS is driven by differences in the epigenomic states of primary pretreatment tumor.

## Genomic landscape of synovial sarcoma

To comprehensively analyze the epigenome of SyS we profiled 52 fresh frozen human SyS tumors following the International Human Epigenome Consortium^29^ (IHEC) standards including total RNA-seq, whole genome bisulfite, and a reference panel of 6 histone modifications supplemented with H3K36me2 and H2AK119Ub1 in a subset of the samples. Whole genome somatic variant analysis was performed on 28 matched tumour/normal pairs. A schematic overview of the study design is represented in **Fig. 1a** and associated clinical metadata presented in **Supplemental Table 1**.

**Figure 1.**
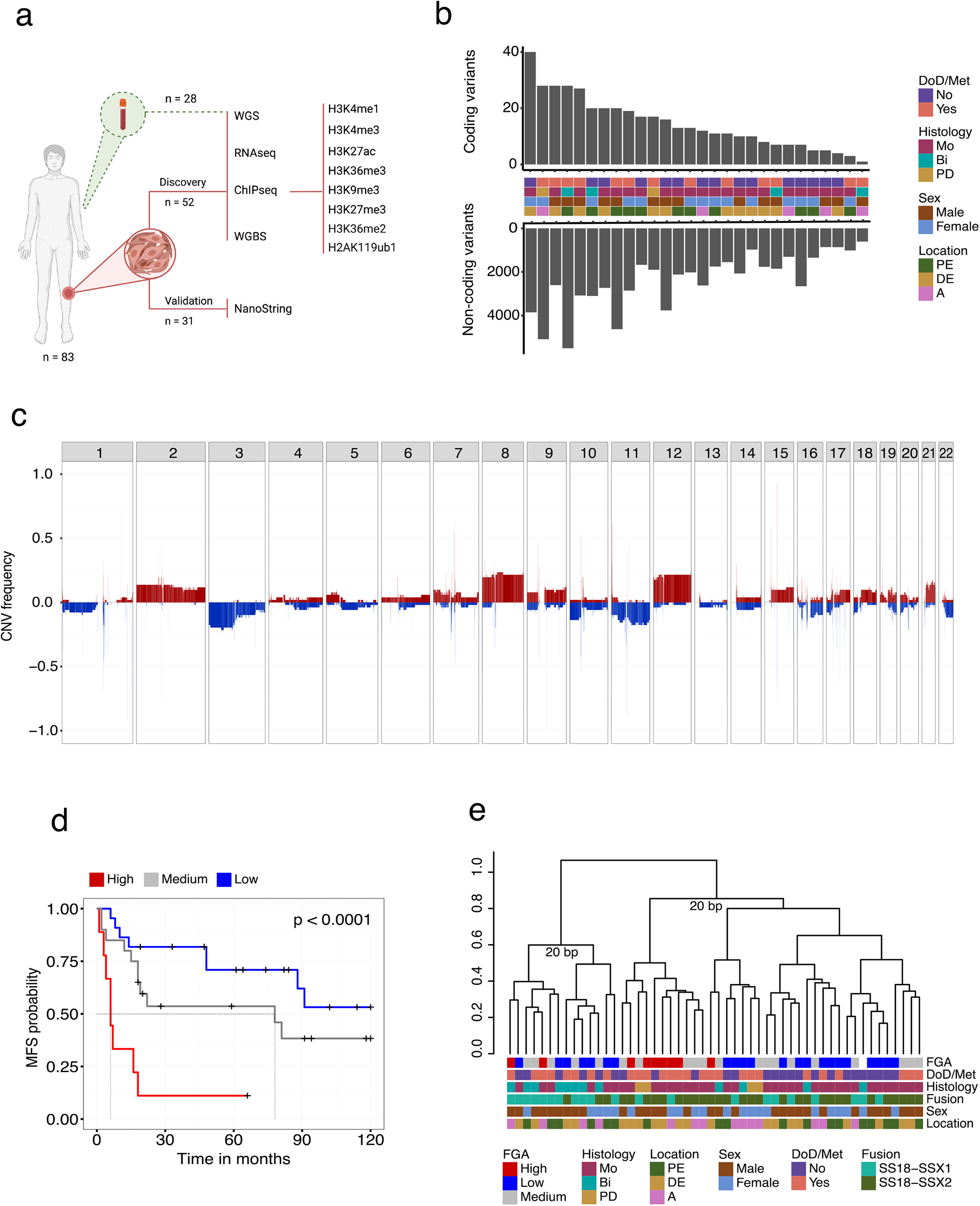
Whole genome landscape of Synovial sarcoma. (**a**) Schematic sample overview. (**b**) Upper;. Barplot showing the number of amino acid altering mutations per sample; Lower; Barplot showing the number of non-coding SNVs and indels per sample. (**c**) Whole genome copy number frequency plot based on CNV calls from WGBS data showing the proportion of samples harboring either gains (red) or losses (blue) across autosomes. (**d**) Kaplan–Meier (KM) survival analysis of the fraction of genome altered (FGA) groups. (**e**) Dendrogram showing unsupervised hierarchical clustering of the top 15% most variably expressed genes (distance = 1 - spearman correlation, ward.D2 clustering method). Abbreviations: DoD, dead of disease; Met, metastasis; Bi, biphasic; Mo, monophasic; PD, poorly differentiated; A, axial; DE, distal extremity; PE, proximal extremity; BP, bootstrap probability.

Consistent with previous studies using targeted gene panels^30,31^ whole genome analysis revealed that SyS genomes are characterized by low numbers of coding SNVs and indels (median, 12.5; range, 1–40) dominated by missense mutations (**Fig. 1b**). No recurrent coding SNVs were detected in the cohort and 9 genes were found to be mutated in more than one tumor genome (**Supplemental Fig. 1a**, **Supplemental Table 2**). Similarly, non-coding SNVs and indels were sparse (median, 2048; range, 609-5504) with no recurrent events nor enrichment in enhancer states. The most common single base substitution mutational signatures (SBS1, SBS5, SBS8; SBS40, SBS89; **Supplemental Fig. 1b**) in SyS have either clock-like (aging associated) or unknown etiologies^32^.

Somatic copy number variants (CNVs) were more common and primarily impacted whole chromosome arms, with amplification of chromosome 8, 12 and loss of 3p being the most common (**Fig. 1c, Supplementary Fig. 1c**). As a group, patient tumours harbouring recurrent CNVs did not display significantly worse prognosis (**Supplementary Fig. 2a-b**), however tumours with a high fraction of genome altered (FGA) were significantly enriched (log-rank test, p < 0.0001) for metastatic disease (**Fig. 1d**). No recurrent high-level amplifications (> 4 copies) were observed and only one recurrent (2/52) homozygous deletion was identified. Taken together, whole genome analysis of SyS confirms a scarcity of recurrent coding SNVs. This observation extends to non-coding regions, and confirms enrichment of recurrent CNV events in cases that metastasize^33^.

## Enhancer signatures define synovial sarcoma molecular subtypes

Next, we explored the epigenomic states of SyS tumors with the aim of identifying molecular subtypes^34–36^. Unsupervised clustering of normalized protein coding gene expression revealed two unstable groups (bootstrap <70)^37^ that were not associated with the available clinical features (**Fig. 1e**), supporting previous work suggesting that SyS tumors do not form distinct clinically relevant subgroups based on transcriptome data^25^.

To explore whether histone modification or DNA methylation could classify SyS we performed unsupervised hierarchical clustering based on genome wide density of the 8 histone modifications, profiled individually, and genome wide fractional methylation calls. Fractional DNA methylation and repressive histone modifications failed to generate stable subgroups; however, two stable and largely consistent subgroups were identified for the active marks H3K4me1, H3K4me3 and H3K27ac (**Supplemental Fig. 2**). Although these subgroups did not align with clinical features and did not show significant correlations with patient outcome.

We focused on active enhancer (H3K27ac) subgroups given their stability and the reported impact of SS18::SSX on the enhancer landscape in SyS cell lines^25^ (**Fig. 2a**). To enable comparative analysis, we first identified a set of group specific enhancers (log2FC > 1; FDR < 0.05). Ranking the tumors based on the difference in normalized read coverage in group specific enhancers revealed a continuous distribution of signal intensity (**Fig. 2b**). To explore the functional significance of the distribution we selected samples from the upper (Q1) and lower (Q4) quartiles of the distribution. Plotting Q1 and Q4 specific enhancers based on their distance from the closest transcription start site (TSS) revealed a significant difference in their genomic distribution, with Q1 peaks enriched at distal regions compared to the Q4 peaks located in closer proximity to TSSs (**Fig. 2c**). In agreement with the differences in genomic distribution, motif enrichment analysis of the subgroup specific peaks identified differences in transcription factor family motifs between the groups^38^, suggesting differences in cBAF and GBAF activity (**Fig. 2d**).

**Figure 2.**
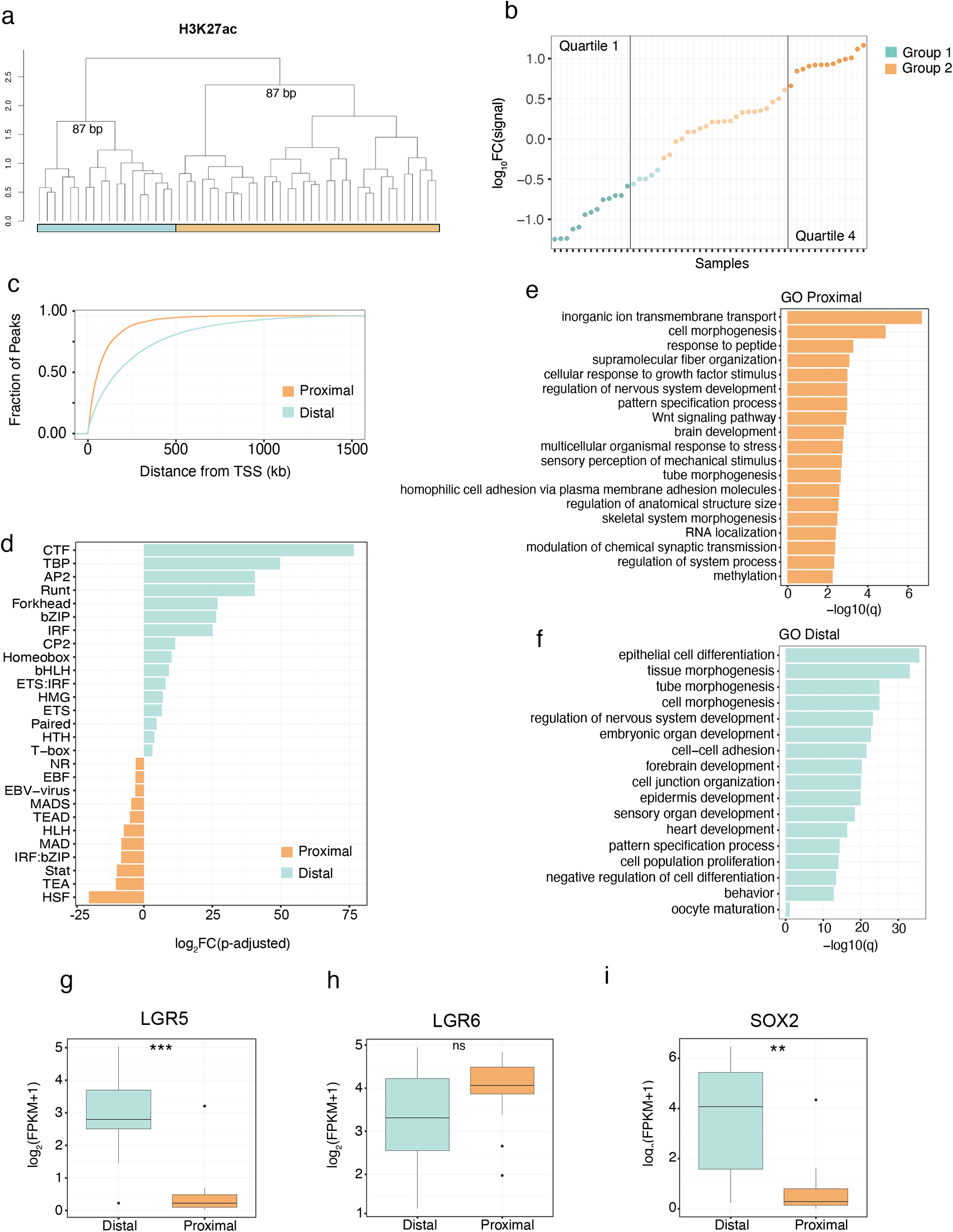
Distinct H3K27ac marker enhancer groups. (**a**) Unsupervised hierarchal clustering of the top 1 % most variable 500bp bins of genome wide H3K27ac signal (distance = 1 - spearman correlation, ward.D2 clustering method). (**b**) Log10(signal intensity) of group 1 specific enhancer regions – log10(signal intensity) of group 2 specific enhancer regions. **(c)** Fraction of the group-specific peaks plotted against the distance to the transcription start site of genes (TSS) for the upper (Q1) and lower (Q4) quartiles of the distribution. (**d**) Fold change difference of transcription factor family motifs enriched at either the distal or proximal group specific enhancers. Gene Ontology enrichment analysis of differentially expressed genes between the (**e**) proximal and (**f**) distal group. Difference in expression between the distal and proximal groups of stem and epithelial markers including (**g**) *LGR5*, (**h**) *LGR6* and (**i**) *SOX2* . ** indicates *P*-value < 0.01, *** indicates *P*-value < 0.001 for a Welch two-sample t test. Abbreviations: BP, bootstrap probability.

Considering the established connection between SS18::SSX and the destruction of the cBAF complex^14^, we posited that variations in the enhancer landscape across primary SyS tumors might be influenced by differing levels of SS18::SSX. However, we did not observe a significant group specific difference in the expression level of *SS18*::*SSX*, including isoforms (**Supplemental Fig. 3a**, b). Similarly, western blot analyses using a fusion-specific antibody did not establish a correlation between the levels of SS18::SSX protein and enhancer groups (**Supplemental Fig. 3c and d**). Additionally, no association between the levels of SS18::SSX, AIRID1A, BRG1, PBRM1, and BRD9 and the enhancer-defined groups were identified by immunohistochemistry (**Supplemental Fig. 3e**).

Having established that the enhancer groups were not a direct reflection of differences in cBAF, GBAF and SS18:SSX levels, we next examined whether these groups are reflective of different tumor trajectories. Supervised differential gene expression analysis was performed between tumors categorized with either distal or proximal enhancers and gene ontology (GO) enrichment analysis was performed on the resulting gene list. Distal enhancer enriched tumors were found to be enriched in epithelial differentiation and keratinization pathways while the proximal enhancer tumor group was enriched in Wnt signaling pathways, among others (**Fig. 2e and f**). Both groups shared pathways involved in neuron and neural tube development. Interestingly, *LRG5* and *LRG6*, which mark distinct mesenchymal lineages in the mouse^39^, were found to be differentially expressed across the enhancer groups (**Fig. 2g and h**). Furthermore *SOX2*, implicated in adult self-renewing epithelia, was specifically expressed in *LRG5* distal enhancer group (**Fig. 2i**). Taken together, these results suggest that SyS tumors exist in a continuum between two enhancer sub-groups that are consistent with distinct mesenchymal progenitor cell types suggesting the possibility of multiple cells of origin.

## Systemic disruption to polycomb mediated repression in primary synovial sarcoma

Recent *in vitro* studies have demonstrated the ability of the C-terminal SSX domain to interact with the nucleosome acidic patch, with a preference for H2AK119Ub1 modified nucleosomes^20,21^. This has led to a model of the aberrant recruitment of GBAF via SS18::SSX to H2AK119Ub1-modified nucleosomes, eviction of H3K27me3, and subsequent gene activation. To explore these relationships *in vivo* we investigated the relationship of H2AK119Ub1 to other histone marks. We first examined the co-occupancy of the 7 histone marks with H2AK119Ub1 in a non-SyS specimen, a *BCOR*-rearranged sarcoma included in our study, and confirmed an expected strong co-association with H3K27me3^40^ (**Fig. 3a**). Strikingly this relationship is disrupted in SyS tumor genomes which instead display the highest co-occupancy between H2AK119Ub1 and the active marks H3K4me3, H3K4me1 and H3K27ac (**Fig. 3b**). To test the dependence of the H2AK119Ub1 overlap and active marks on SS18::SSX we performed ChIP-seq on the synovial sarcoma cell line SYO1 before and after SS18::SSX siRNA mediated knockdown (**methods; Supplemental Fig. 3f**). Co-occupancy of H2AK119Ub1 and active marks was also apparent in the SYO1 cell line and was reduced following SS18::SSX2 knockdown (**Fig 3c**). We further confirmed the SS18::SSX dependency of active marks and H2AK119Ub1 in a primary mouse cell line (C3H10T1/2) stably expressing SS18::SSX1^24^ (**Supplemental Fig. 3g**) and in a novel Hic1 SyS mouse model^41^ (**Fig. 3d**). Taken together, our data supports a model where SS18::SSX disrupts canonical PRC1-PRC2 complex mediated repression, driving a loss of H3K27me3 but retention of H2AK119Ub1 during the subsequent acquisition or retention of active histone marks.

**Figure 3.**
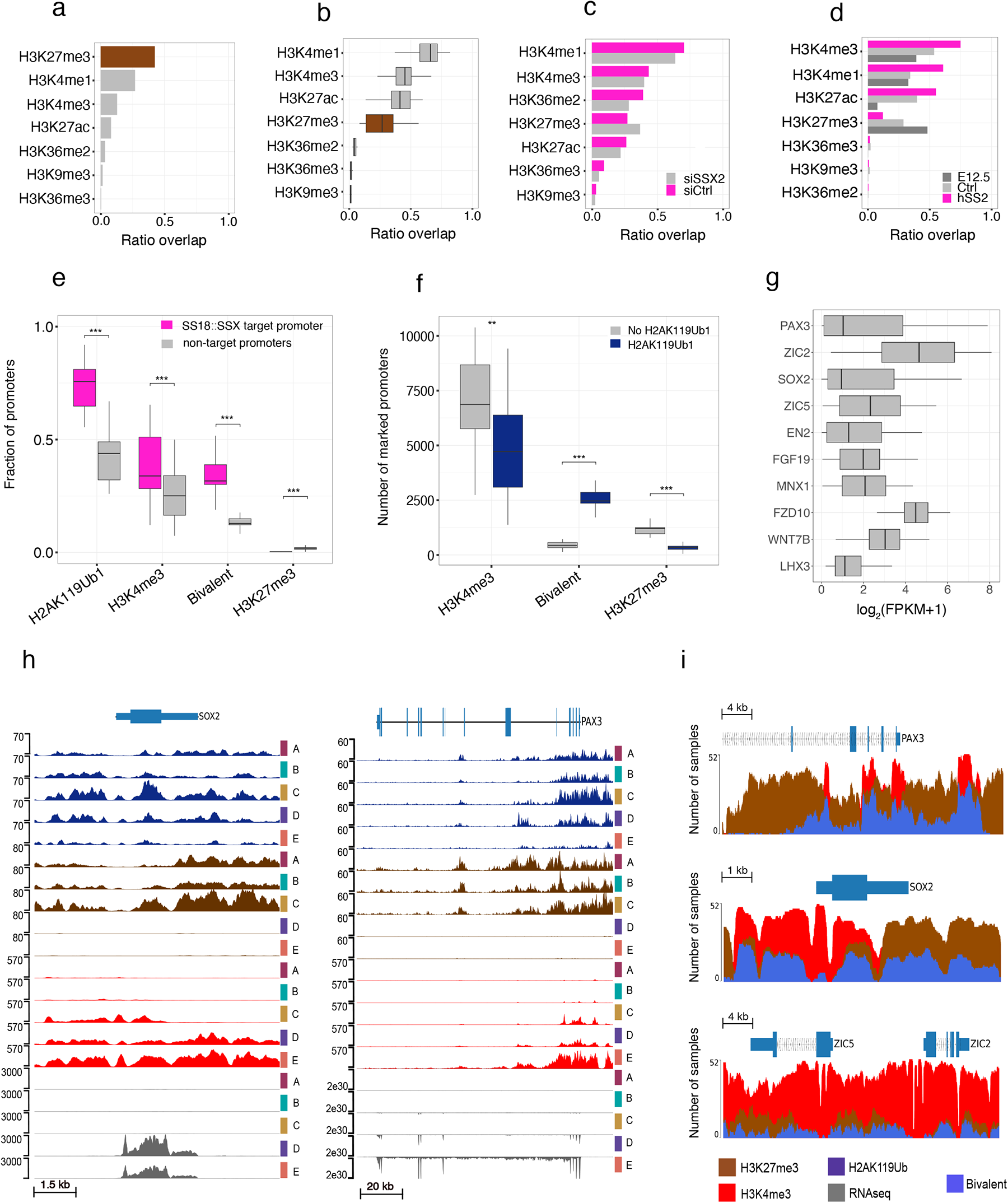
H2AK119Ub1 and variable target gene activation. Fraction of genome wide H2AK119Ub1 occupancy overlapping other histone marks in (**a**) a non-SyS samples, (**b**) SyS tumors, (**c**) the SYO1 cell line treated with either siRNA against SSX2 (siSSX2) or scramble control (siCtrl), (**d**) HIC^+^ driven mouse tumor (hSS2), adult HIC^+^ MSCs (Ctrl) or embryonic MSCs (E12.5). (**e**) Fraction of protein coding promoters that are marked by H2AK119Ub1 and are either SS18::SSX targets or not. Promoters are then further subdivided into H3K4me3 (H2AK119Ub1 and H3K4me3 without H3K27me3), bivalent (H2AK119Ub1 and H3K4me3 and H3K27me3) or H3K27me3 (H2AK119Ub1 and H3K27me3 without H3K4me3). (**f**) The number of protein coding promoters marked by H3K4me3 (H3K4me3 without H3K27me3), bivalent (H3K4me3 and H3K27me3) or H3K27me3 (H3K27me3 without H3K4me3). Promoters are then further subdivided into being marked by H2AK119Ub1 or not. (**g**) Gene expression values of signature synovial sarcoma genes ordered by variance. (**h**) ChIP-seq signal enrichment and gene expression tracks for five SyS primary tumors at the *SOX2* and *PAX3* locus. H2AK119Ub1 (dark blue), H3K27me3 (brown), H3K4me3 (red) and gene expression (grey) are displayed. (**i**) Visual representation of peak calls for H3K4me3 (red), H3K27me3 (brown) or bivalent (blue) regions around the *SOX2*, *PAX3, ZIC2* and *ZIC5* genes. The amplitude represents the number of samples that have a called peak in that region. ** indicates *P*-value < 0.01, *** indicates *P*-value < 0.001 for a Welch two-sample t test.

Next, we focused specifically on SS18::SSX target promoters, determined by SS18::SSX ChIP-seq in SyS cell lines^25^, and found as expected that H2AK119Ub1 was enriched at promoters bound by SS18::SSX compared to all other protein coding promoters (**Fig. 3e**). Consistent with an altered relationship between H2AK119Ub1 and active marks we found that a significant fraction of H3K4me3 marked promoters were co-marked by H2AK119Ub1. Conversely, we observed a reduction in the proportion of promoters marked by H3K27me3 and H2AK119Ub1 at SS18::SSX targets. We next looked at the relationship between H2AK119Ub1 and bivalent promoters (co-marked by H3K4me3 and H3K27me3) and observed a significant increase (t-test, p=5.4e-10) at SS18::SSX target sites compared to all other promoters (**Fig. 3e, f**). In agreement with the observed heterogeneity of active and repressive chromatin states at SS18::SSX target genes, we observed highly variable expression of reported SyS sentinel genes (**Fig. 3g**). Indeed, manual inspection of the genes, *SOX2*, *PAX, ZIC2* and *ZIC5*, revealed a remarkable heterogeneity between active and repressive marks across the cohort of primary tumors (**Fig. 3h and i**). This observation is in agreement with the heterogeneous SS18::SSX-mediated gene activation observed in SyS cell lines^17^.

## Increased bivalency is associated with poor outcome in synovial sarcoma

Having established a high degree of heterogeneity of chromatin states at sentinel genes across the cohort we asked whether this heterogeneity was reflective of tumor state. We focused on bivalent promoters and noted that the number of bivalent promoters was highly variable across the cohort and, intriguingly, that a reduced number of bivalently marked promoters appeared to be associated with favorable outcome (**Fig. 4a**). To explore this further, we split the distribution into two groups and classified the tumors into one of the two groups, bivalency high (bivH) or bivalency low (bivL), based on their number of marked promoters. The Cox proportional-hazards model^42^ was used to test the association between metastasis free survival (MFS) time and the following predictor variables: sex, fusion variant, anatomic location, bivalency group and bivalency counts. The only variables with significant impact on MFS time were bivalency group and the bivalency count itself (**Supplementary table 3**) and the grouping alone was strongly prognostic (**Fig. 4b**).

**Figure 4.**
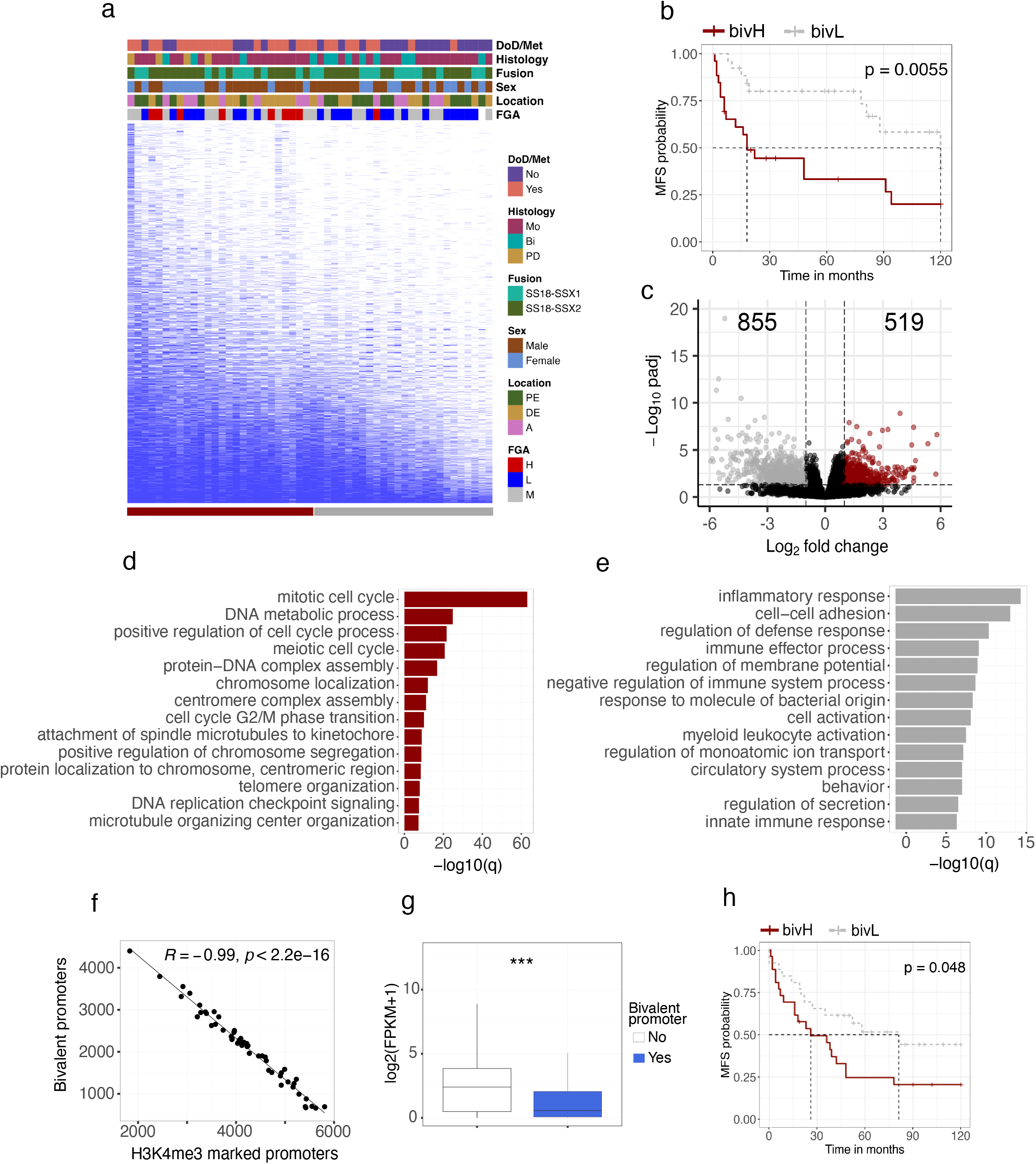
Bivalency is a prognostic marker for synovial sarcoma. (**a**) Binary heatmap over bivalently marked promoters (rows) in synovial sarcoma samples (columns) ordered by the total number of marked promoters. **(b)** Kaplan–Meier (KM) analysis of metastasis-free survival (MFS) in bivalency high (bivH) and bivalency low (bivL) groups. Statistically significant difference between the curves was calculated using the log-rank test. **(c)** Volcano plot displaying the number of differentially expressed genes between the upper and lower quartiles of the bivH and bivL groups. Gene Ontology enrichment analysis of genes found to be (**d**) upregulated or (**e**) downregulated in the bivH quartile. (**f**) Spearman correlation between the number of bivalently marked promoters and the number of promoters marked by H3K4me3 only. (**g**) Box plot showing the gene expression values for promoters being either bivalently marked or not. (**h**) KM analysis of MFS in the validation set, comparing the difference in survival between tumors predicted to be bivH or bivL. Statistically significant difference between the curves was calculated using the log-rank test. *** indicates *P*-value < 0.001 for a Welch two-sample t test. Abbreviations: DoD, dead of disease; Met, metastasis; Bi, biphasic; Mo, monophasic; PD, poorly differentiated; A, axial; DE, distal extremity; PE, proximal extremity.

To explore the functional significance of bivalency levels we performed differential expression analysis between the groups and identified 1374 differentially expressed genes (FDR < 0.05) (**Fig. 4c and Supplementary table 4**). Consistent with the prognostic relationships we observed significant enrichment of GO terms associated with cellular proliferation in the gene set upregulated in the bivH group including numerous terms associated with cell cycle and DNA replication (**Fig. 4d**). In addition to these terms, we identified several oncogenes including c*MYC*, whose expression was strongly correlated with the number of bivalently marked promoters (**Supplemental Fig. 4a**).

GO analysis of genes upregulated in bivL tumors (down in bivH) revealed numerous terms related to immune response (**Fig. 4e**). Although SyS is an immune cold tumor type with low levels of immune cell infiltrates^43^ single cell RNA-seq of SyS tumors identified a malignant cellular state associated with poor prognosis and immune evasion^44^. In agreement with our results, analysis of these SyS cell type specific signatures^44^ identified significantly higher expression of several immune cell type signatures, particularly mastocytes, within the bivL group (**Supplemental Fig. 4b**). This observation was further supported by significant differences in estimated immune cell factions from transcriptome deconvolution analysis (**Supplemental Fig. 4c**). In addition, enrichment of the malignant programs, “cell cycle” and “core oncogenic program” (**Supplemental Fig. 4d and e**), which have been shown to be associated with poor prognosis and to anti-correlate with the level of immune infiltrates^44^, was significantly correlated with bivalency. Taken together, these results suggest that the unfavorable outcome observed in the bivH tumors is linked to an increased expression of cell cycle associated genes, including *MYC*, in combination with downregulation of the immune response.

## Bivalency is prognostic in synovial sarcoma

We noted a striking directionality to the loss of bivalency across the cohort with a strong correlation (R=0.99) to the number of promoters that resolve to an active state marked only by H3K4me3 (**Fig. 4f**) while no correlation between loss of bivalency and gain of promoters marked by H3K27me3 was observed (**Supplemental Fig. 4f**). In addition, genes whose promoters lose bivalency increase in gene expression (**Fig. 4g**) supporting a model of a strong directional loss of bivalent states to an active state across the cohort. Accordingly, we reasoned that gene expression levels could be used as a proxy for bivalency in a validation cohort.

To design a clinically relevant SyS bivalency test, we opted to use the NanoString nCounter System to directly quantify mRNA-expression patterns, as it has been shown to be well suited for handling clinical specimens in a rapid and cost-effective manner^45^ and is in clinical use for sarcoma diagnosis^46,47^. A NanoString CodeSet designed to target 50 marker genes (that switch from a bivalent to an expressed active state) was constructed and used to analyse the mRNA-expression levels for all samples across both cohorts **(Supplemental Table 5**). The gene-gene correlation was high (mean 0.89, spearman) when the expression of the marker genes was compared between RNA-seq and NanoString in the discovery cohort (**Supplemental Fig. 4g**).

To weigh the individual expression values, a random forest model was trained on the 31 samples in the discovery cohort. The resulting model was assessed using receiver operating characteristic (ROC) analysis which showed that the expression signature was highly predictive of bivalency group (AUC = 0.96) (**Supplemental Fig. 4h**). The model was then used to predict the bivalency status of the validation cohort (N=51: **Supplemental Table 1)**, classifying 23 samples as bivH and 28 samples as bivL, and survival analysis confirmed a significant difference in survival between the predicted bivalency groups (**Fig. 4h**). Taken together, these results support a model where poor outcome is associated with increased levels of bivalency and provides the basis for a clinically applicable test.

BivL tumors demonstrated particularly low levels of genomic alterations (**Supplemental Fig. 4i**). These tumors were also smaller in size and harbored fewer SNVs, but interestingly had significantly higher expression of fusion target genes (**Supplemental Fig. 4 j,k and l**), suggestive of a state driven by SS18::SSX. Conversely, this also suggests that elevated genomic instability may reduce the dependence on SS18::SSX, diminishing the requirement of conversion of bivalent promoters into an active state.

## Synovial sarcoma genomes show distinct H3K4me3 patterns

Studies in SyS cell lines, mouse models and organoids have consistently observed the formation of unusually broad BAF domains, associated with the loss of H3K27me3 and gain of active histone marks, as a hallmark of the SS18::SSX fusion protein^14,24,25^. To investigate whether such alterations would translate into observable changes in genome wide occupancy of histone marks in primary human tumors, we assessed the genomic occupancy patterns of histone modifications of SyS in the context of the IHEC EpiATLAS collection^48^. Unsupervised clustering of promoter associated histone modification promoter occupancy revealed that SyS genomes harbor distinct histone modification landscapes compared to tissues profiled in the EpiATLAS (**Supplemental Fig. 5a, b and c**). Among the core histone marks profiled by IHEC, H3K4me3 was the most distinctive in this analysis demonstrating not only a unique promoter occupancy pattern but also significantly increased density (**Fig. 5a**). Increased density of H3K4me3 at promoters extended genome-wide where its occupancy was found to be significantly increased (pairwise wilcoxon signed-rank test, p < 0.05) in SyS compared to the EpiATLAS compendium (**Fig. 5b, Supplemental Fig 6a-e**). Consistent with broad BAF domains at SS18::SSX target genes, a significant expansion of H3K4me3 was also evident at promoter regions, where SyS displayed the widest domains of any tissue (**Fig. 5c**). These findings support a direct relationship between the observed H3K4me3 expansion and the SS18::SSX recruitment, and we observed a significant (t-test, p = 4e-09) increase in the H3K4me3 domain width at fusion binding sites compared to SS18 sites **(Supplemental Fig 6f).**

**Figure 5.**
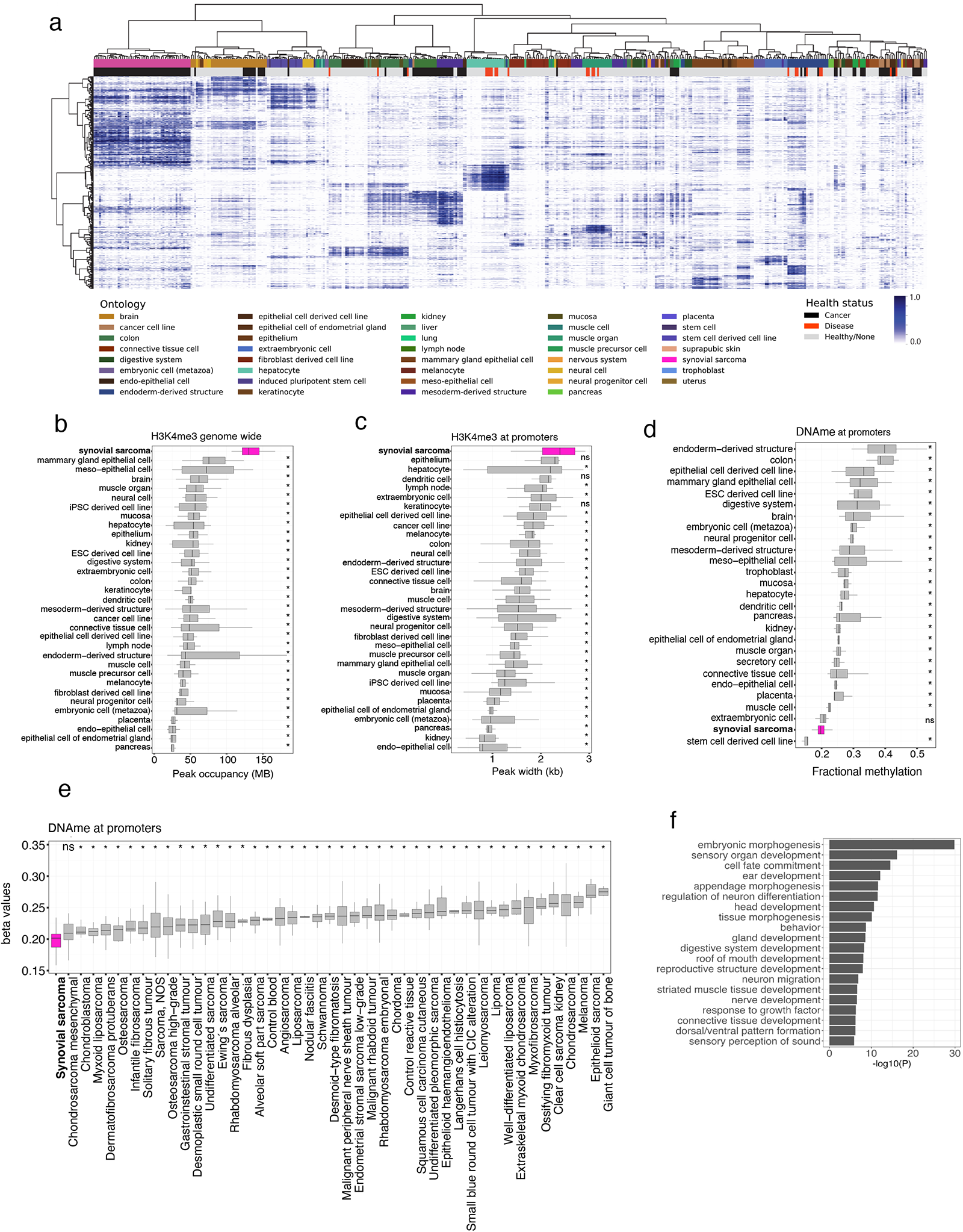
Synovial tumors harbor broad H3K4me3 domains. (**a**) Unsupervised hierarchal clustering (distance = 1 - spearman correlation, ward.D2 clustering method) of H3K4me3 in primary SyS (pink) and other normal and diseased tissue from the International Human Epigenome Consortium (IHEC). Heat is the proportion of promoter regions marked by H3K4me3 peaks. Box plots showing **(b)** the genomic occupancy of H3K4me3, **(c)** mean peak width of H3K4me3 at promoter regions and **(d)** fractional methylation of promoters in SyS and other normal and diseased tissue from IHEC. **(e)** Mean methylation (beta values) of promoter regions in SyS and other sarcoma subtypes. **(f)** Gene Ontology analysis of genes whose promoters are overlap differentially hypomethylated regions (DMRs) in SyS compared to other sarcoma subtypes. * indicates *P*-value < 0.05 using pairwise Wilcoxon signed-rank test.

## Synovial sarcoma genomes are characterized by DNA hypomethylation

We hypothesized that the broad H3K4me3 signature in SyS would translate into a DNA hypomethylation phenotype given known antagonisms between H3K4me3 and DNA methylation^49^. As expected, we observed strong anti-correlation between H3K4me3 and DNA methylation at promoter regions (-0.95, spearman). Consistent with increased H3K4me3 occupancy, in comparison to the EpiATLAS collection, SyS genomes demonstrated the lowest mean promoter methylation, apart from stem cell derived cell lines (**Fig. 5d**). The same hypomethylated phenotype was not observed genome wide, in gene bodies or intergenic regions (**Supplemental Fig. 6g, h and i**)

To further explore the specificity of SyS DNA promoter hypomethylation, we leveraged methylation array data from over 1,500 sarcoma samples spanning over more than 50 histological subtypes^50^. Strikingly, SyS displayed the lowest mean methylation level of any sarcoma confirming this feature is distinctive to SyS (**Fig. 5e and Supplemental Fig. 7a**). Differential methylation analysis revealed that 90% of differentially methylated probes (DMPs) and 97% of regions (DMRs) were hypomethylated in SyS compared to other sarcomas subtypes. Hypomethylated SyS DMRs span promoter regions of several genes reported to be direct targets of SS18::SSX and/or key drivers of synovial sarcomagenesis, including *SOX2, PAX3, SIM2, TWIST1* and *PDGFRA*^18,25,51,52^. The majority of hypomethylated DMRs (74%) span promoter regions and GO analysis of the associated genes revealed a significant enrichment of embryonic and neural pathways suggestive of a remnant of an epigenetic signature associated with a putative cell-of-origin (**Fig. 5g**). Motif enrichment analysis of the hypomethylated DMRs revealed an almost exclusive enrichment of homeobox transcription factors **(Supplemental Fig. 7b)**. KDM2B, a genetic dependence in SyS, that specifically recognizes non-methylated CpGs to recruit the non-canonical polycomb repressive complex (ncPRC1.1) to catalyze H2AK119Ub1 deposition was not identified among the most significantly enriched motifs.

Promoters harbouring CGIs were more strongly hypomethylated than promoters lacking CGIs in SyS compared to other tissues (**Supplemental Fig. 7c and d**) and thus we posited that the observed SyS hypomethylation was due in part to a spreading of CGI hypomethylation into the surrounding CGI shores. Indeed, when focusing on CGI shores, SyS displayed the lowest methylation level apart from embryonic stem cells (**Fig. 6a**) and a gradual decrease in methylation towards the CGIs (**Supplemental Fig. 7e**).

**Figure 6.**
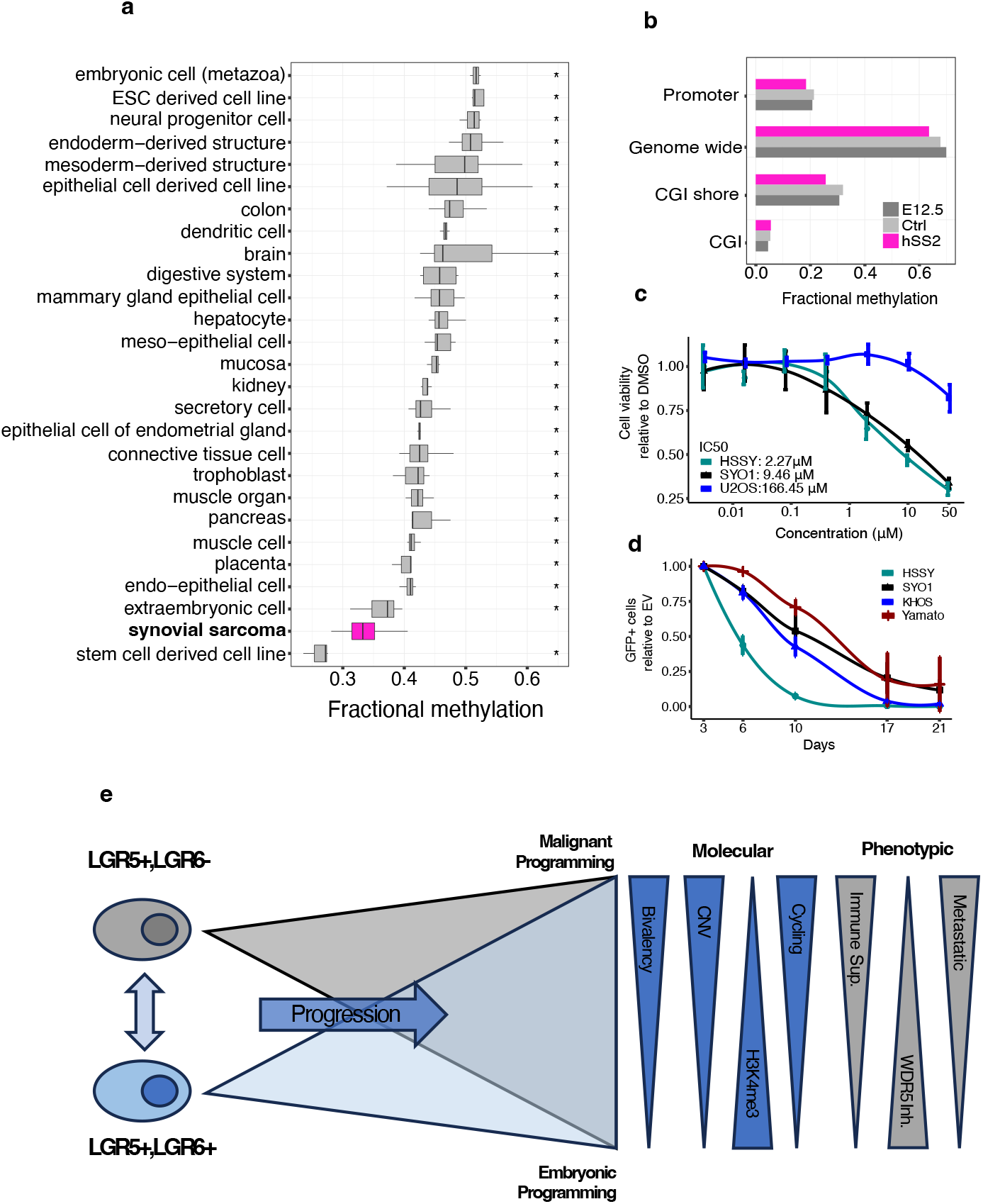
CGI hypomethylation associated with cell of origin and elevated H3K4me3 provides a potential therapeutic vulnerability. (**a**) Fractional methylation of SyS and a collection of IHEC tissues in CGI shores**. (b)** Fractional methylation of genome wide, promoters, CGIs and CGI shores in the hSS2, Ctrl and E12.5 mouse samples. (**c)** Cell viability assay in SyS (HYSSII and SYO1) and Osteosarcoma (U2OS) cells upon 7-day treatment with the WDR5 inhibitor OICR-9429. **(d)** Cell competition assay performed in the SyS lines (HSSYII, SYO1, and Yamato) and Osteosarcoma (KHOS) cells transduced with an empty sgRNA as control or with guides targeting WDR5. **(e)** Proposed summative model. * indicates *P*-value < 0.05 using pairwise Wilcoxon signed-rank test.

Given their role in maintaining DNA methylation homeostasis at CGI boundaries we compared the expression levels of DNMT and the TET gene family^53^ and found *TET1* to be highly expressed in SyS compares to the IHEC tissue compendium (**Supplemental Fig. 7f**). We confirmed high levels of *TET1* expression in a SyS mouse model (**Supplemental Fig. 7g**), providing an opportunity to directly test whether the *TET1* overexpression and the associated hypomethylated phenotype was acquired during tumorigenesis or was a reflection of an embryonic cell-of-origin. We performed whole genome bisulfite analysis of the HIC1+ population at E12 and in the presence and absence of human SS18::SSX2^41^. DNA methylation levels mirrored that seen in the primary human tumours including extreme hypomethylation at promoters and CGI shores (**Fig. 6c**). However, a hypomethylated phenotype was also largely present in the matched normal cells with a modest decrease in methylation observed following tumour induction (**Fig. 6c**). We further reasoned that if *TET1* mediated hypomethylation was essential for SyS we should observe sensitivity to its inhibition. However, neither 2-Hydroxyglutarate treatment^54^ nor *TET1* CRISPR knockout had a selective impact on SyS cell lines (SYO1, HSSYII, Yamato) compared with osteosarcoma cell lines (U2OS, KHOS) (**Supplemental Fig. 7h and i**). Collectively, these results suggest that while DNA hypomethylation is a defining characteristic of SyS , and likely contributes to tumorigenesis as a requirement for KDM2B^18^ recruitment, it is a reflection of cell-of-origin rather than a consequence of SS18::SSX expression.

## H3K4me3 is an epigenetic vulnerability in synovial sarcoma

The uniquely broad H3K4me3 regions in human SyS tumors, together with the demonstrated ability of SS18::SSX to generate broad active regions^24,25^ prompted us to explore whether H3K4me3 inhibition could be a specific dependency in SyS. Interrogation of genome-scale RNAi viability screens across a diverse range of cancer cell lines (DepMap) identified WDR5 depletion as a selective vulnerability in SyS (**Supplemental Fig. 8a**). WDR5 is core member of the COMPASS and MLL family of complexes and is essential for the enzymatic activity of H3K4me3-specific histone methyltransferase complexes^55^. The inhibition of WDR5 has emerged as a promising therapeutic target in several cancers including acute myeloid leukemia, glioblastoma and bladder cancer^56–58^ . We confirmed that *WDR5,* and the other COMPASS and MLL family members, were consistently expressed in SyS tumors **(Supplemental Fig. 8b)**. Then, to assess the impact of WDR5 inhibition on SyS, we treated cell lines with a WDR5 inhibitor (OICR-9429) and directly knocked it out using a CRISPR base editing. SyS cell lines demonstrated sensitivity to both chemical and genetic inhibition of WDR5 inhibition suggesting that targeting H3K4me3 may be a therapeutic vulnerability in SyS (**Fig. 6d and e**). In agreement with previous results (DepMap, **Supplemental Fig. 8c**), genetic knock out of *WDR5* resulted in reduced proliferation in both the SyS and control cell lines, highlighting its critical role maintaining H3K4me3 homeostasis. Chemical inhibition of WDR5 showed increased sensitivity in SyS compared to the control line suggesting that SyS may be uniquely sensitivity to H3K4me3 perturbation.

Taken together, comprehensive analysis of the epigenome of primary SyS provides insight into the clinical heterogeneity observed in SyS and suggests novel therapeutic and prognostic avenues for SyS management (**Fig. 6f**).

## Discussion

SyS is driven by a pathognomonic translocation event and yet displays histologic and clinical heterogeneity. Comprehensive genomic analysis reinforces previous targeted efforts demonstrating a paucity of recurrent coding SNVs that extends into non-coding space.

Collectively, this suggests that unlike many cancers, SyS does not typically rely on driver mutations in its evolution, emphasizing the potential importance of other regulatory mechanisms in its pathogenesis. Low level chromosome scale recurrent CNVs are observed, enriched in copy number gains and linked with progression to metastatic disease. Epigenomic states, notably at bivalent promoters, are surprisingly heterogenous in SyS. However, two stable groups, delineated by active enhancers, are observed that do not appear associated with progression but may instead represent remanent cell-of-origin signatures. Proximal and distal enhancers are also frequently found in association with H2AK119Ub1 in a manner dependent on SS18::SSX. This aberrant relationship between active and repressive marks in the chromatin landscape of SyS is consistent with the understood mechanisms of SS18::SSX reprogramming and likely underpins specific oncogenic processes in SyS.

Bivalency across SyS tumors appears as a continuum with a striking directional resolution towards H3K4me3. Consistent with a key role in SS18::SSX driven reprogramming, increased genomic occupancy of H3K4me3 is a defining feature of the SyS epigenome where it occupies broad domains at TSSs. Broad H3K4me3 domains, antagonistic to DNA methylation, are associated with an extreme DNA hypomethylation signature where SyS is the most hypomethylated of any sarcoma. Low bivalency and increased H3K4me3 in association with H3K27ac and H2AK119Ub1 is consistent with increased levels of SS18::SSX reprogramming and is associated with improved outcome. We posit that this state is reflective of increased SS18::SSX dependency/reprogramming and that the epigenetic state, influenced by SS18::SSX, is crucial in determining the disease course and could serve as a biomarker for prognosis.

Temporal and anatomical specificity of SyS suggests its origins are constrained to cellular states that are permissive to SS18::SSX driven epigenomic reprogramming. Enhancer subgroups support a mesenchymal origin but suggest the possibility of multiple potential cell-of-origins. High levels of *TET1* expression, associated with CGI hypomethylation, also suggests remnants of the cell-of-origin, rather than events triggered by SS18::SSX reprogramming. This finding provides insight into the developmental origins of SyS, potentially influencing therapeutic approaches that target these foundational epigenetic settings.

Finally, the inhibition of H3K4me3 is proposed as a novel therapeutic vulnerability in SyS. Given the role of this histone modification in sustaining the malignant phenotype, targeting this pathway may offer a new avenue for treatment, particularly for those tumors exhibiting extensive H3K4me3 domains.

Comprehensive epigenomic profiling of SyS not only underscores the complexity of its genetic and epigenetic landscape but also opens up new avenues for targeted therapies. Further understanding the mechanisms underlying SyS oncogenesis can help in developing more effective and personalized treatment strategies for patients with SyS.

## Acknowledgements

Sources of support: Funding from National Institutes of Health (1U54CA231652-01), Canadian Cancer Society (#705615), Terry Fox Research Institute (#1082), Terry Fox Research Institute Marathon of Hope Cancer Centres Network (MOHCCN) (#3311-01), the Swedish Childhood Cancer Foundation (TJ2020-0010) and Michael Smith Foundation for Health Research (RT-2020-0601).

## Data Availability

Data accessions and associated public metadata are available through https://www.ebi.ac.uk/epirr

## Supplementary Figures

**Supplemental Figure 1.**
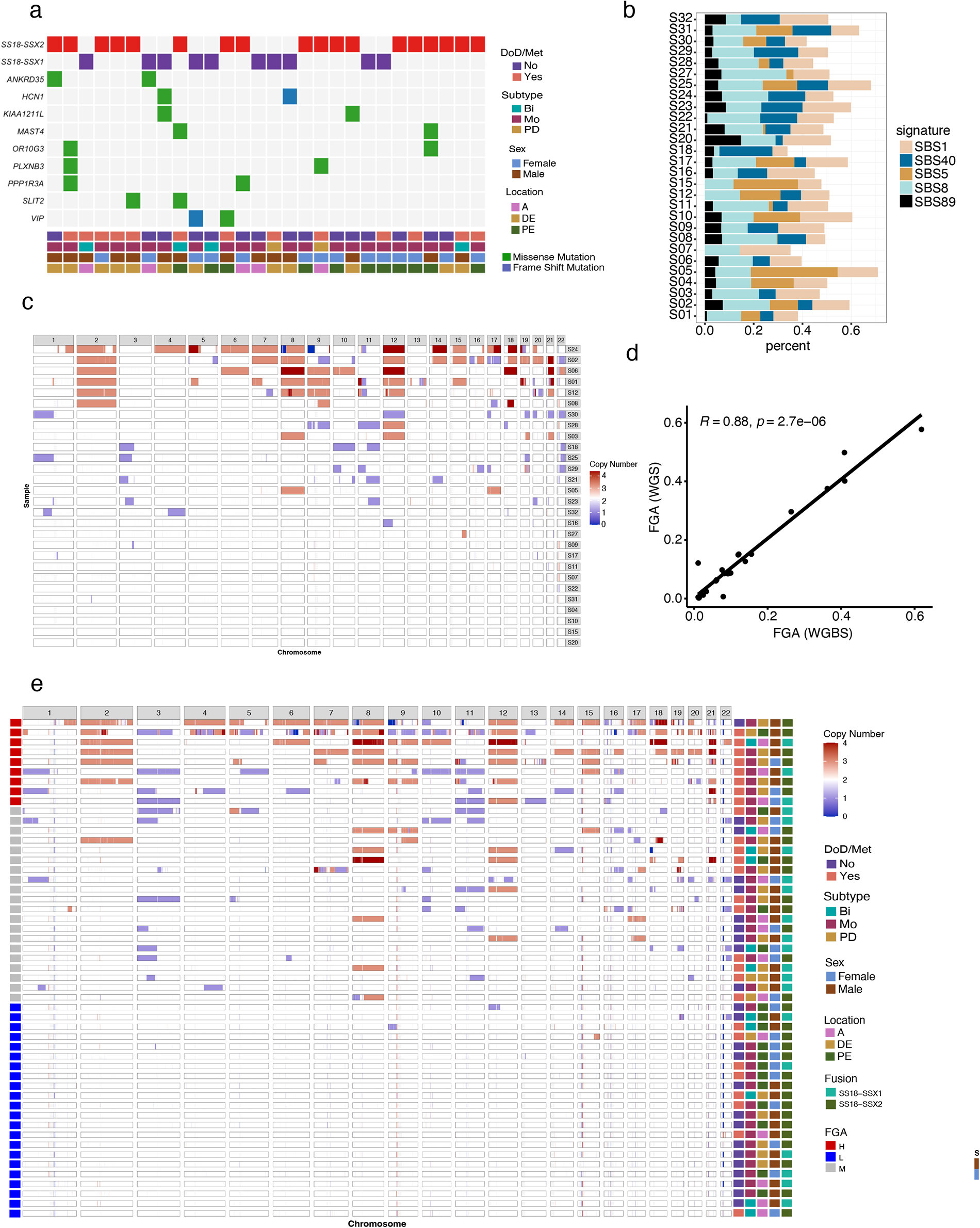
Synovial sarcoma samples do not have recurrent mutations. (**a**) Oncoplot displaying genes affected by missense mutations in more than one sample. (**b**) The frequencies of the five overall most common single base substitution (SBS) signatures per case. (**c**) Whole genome copy number heatmap based on WGS data with samples ordered by the fraction of the genome altered (FGA). **(d)** Spearman correlation between the FGA calculated from WGS and WGBS data. **(e)** Whole genome copy number heatmap base on WGBS data with samples ordered by FGA. Abbreviations: MFS, metastatic free survival; Bi, biphasic; Mo, monophasic; PD, poorly differentiated; A, axial; DE, distal extremity; PE, proximal extremity.

**Supplemental Figure 2.**
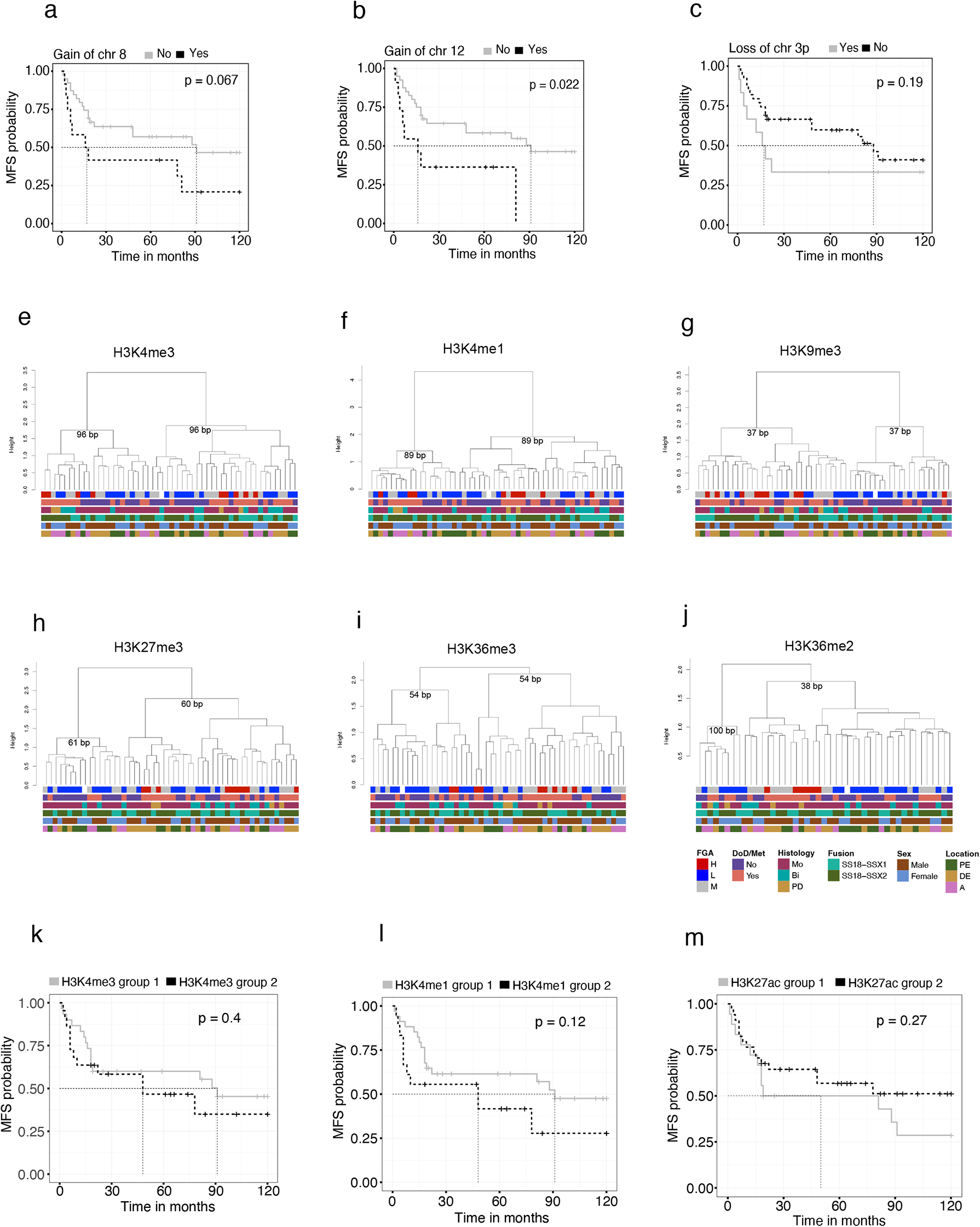
Active histone marks reveal two stable groups but do not have significant correlation with clinical outcome. Kaplan–Meier (KM) survival analysis of samples with or without **(a**) gain of chromosome 8, (**b**) gain of chromosome 12, or (**c**) loss of chromosome arm 3p. Unsupervised hierarchal clustering of the top 1 % most variable 500bp bins (distance = 1 - spearman correlation, ward.D2 clustering method) of genome wide (**d**) fractional methylation, (**e**) H3K4me3, (**f**) H3K4me1, (**g**) H3K9me3, (**h**) H3K27me3, (**i**) H3K36me3 and (**j**) H3K36me2 signal. KM analysis of the stable subgroups (bootstrap value > 70) observed in (**k**) H3K4me3, (**l**) H3K4me1, (**m**) H3K27ac. Abbreviations: DoD, dead of disease; Met, metastasis; Bi, biphasic; Mo, monophasic; PD, poorly differentiated; A, axial; DE, distal extremity; PE, proximal extremity; BP, bootstrap probability.

**Supplemental Figure 3.**
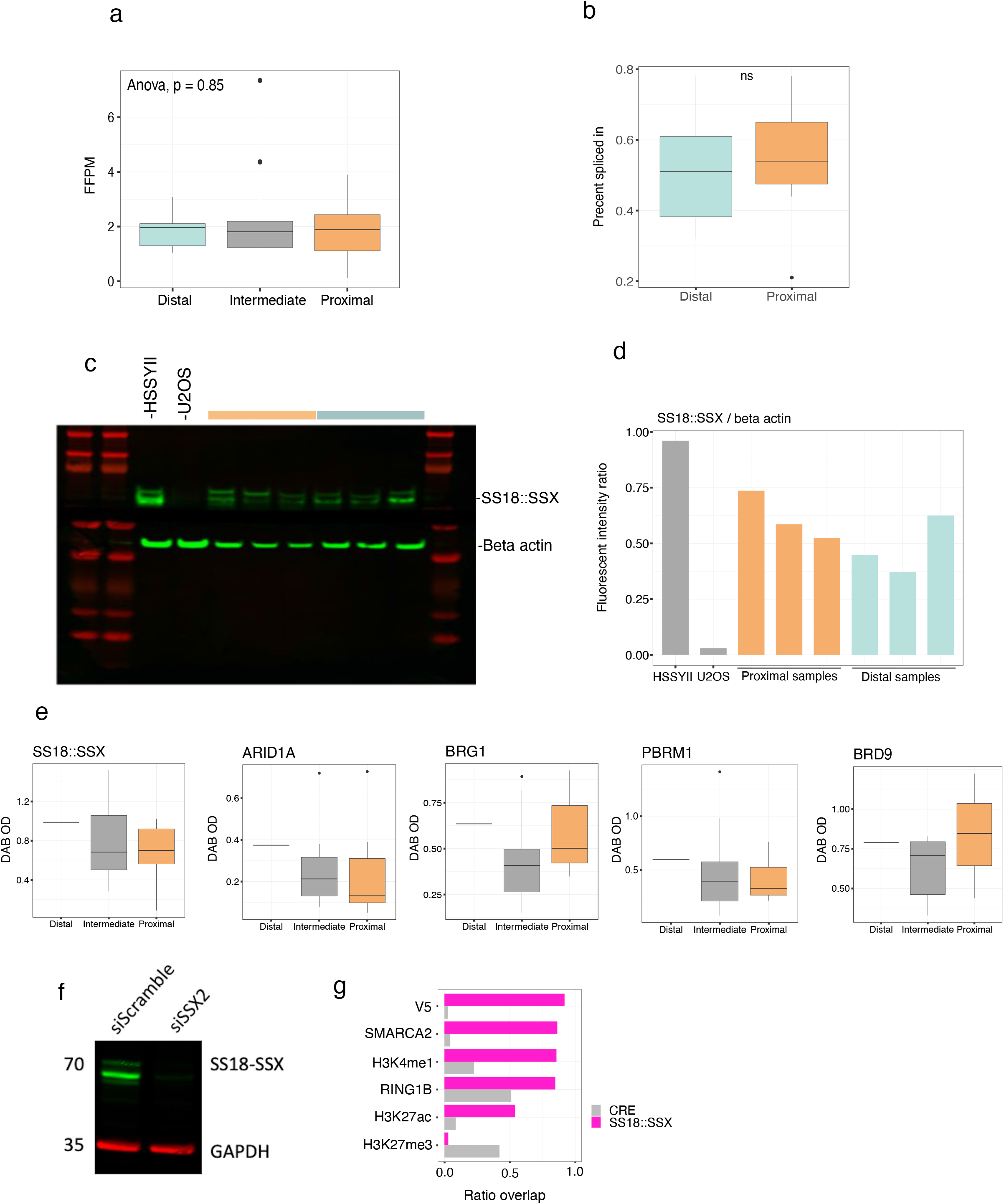
Enhancer defined groups are not explained by SS18::SSX , cBAF, or GBAF levels. (**a**) Number of RNA-seq reads supporting the existence of the fusion per million mapped reads (fusion fragments per million, FFPM). (**b**) Miso analysis of the percentage spliced in (PSI) values for SS18 exon 8 isoforms between proximal and distal groups. (**c**) Western blot showing SS18::SSX and beta actin in HSSY2, U2OS (osteosarcoma cell line), 3 Proximal tumors (orange) and 3 Distal tumors (turquoise). (**d**) Quantified normalized expression of SS18::SSX compared to beta actin from the Western blot in HSSY2, U2OS (osteosarcoma cell line), 3 Proximal tumors (orange) and 3 Distal tumors (turquoise). (**e**) DAB nuclear optical density using antibodies for SS18::SSX, ARID1A, BRG1, PBRM1 and BRD9. (**f**) Western blot to demonstrate siRNA knockdown of SS18::SSX in SYO1 cell lines (5nM, 3 days). (**g**) Fraction of genome wide H2AK119Ub1 occupancy overlapping other histone marks in the C3H cell line expressing SS18::SSX and with SS18::SSX knockdown using CRE.

**Supplemental Figure 4.**
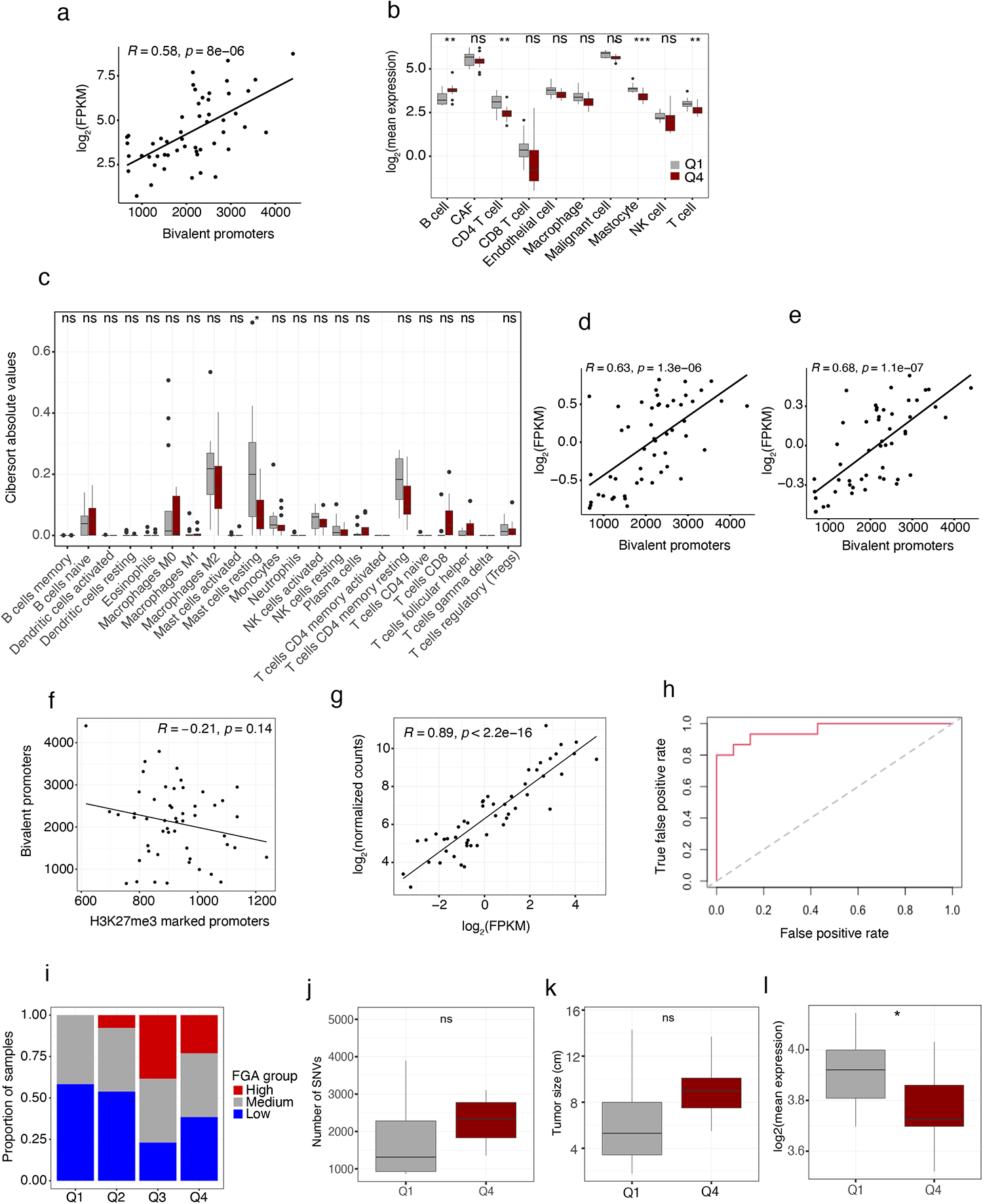
Bivalency level correlates with cell cycle, core oncogenic, and immune cell expression signatures. (**a**) Spearman correlation between the number of bivalently marked promoters and the MYC expression. **(b)** Mean expression of SyS cell type specific signatures in the upper (Q4) and lower (Q1) bivalency quartiles. **(c)** Cibersort absolute values in bivalency high (upper quartile, Q4) and bivalency low (lower quartile, Q1) samples. Spearman correlation between the number of bivalently marked promoters and the mean expression of the SyS cell signature **(d)** “cell cycle” and **(e)** “core oncogenic program”. **(f)** Spearman correlation between the number of bivalently marked promoters and promoters marked with H3K27me3 only. **(g)** Gene-gene correlation between the expression of the marker genes measured by RNA-seq and NanoString in the discovery cohort. **(h)** Receiver-operating characteristic (ROC) curve demonstrating that the NanoString signature was highly predictive of the bivalency groups in the discovery cohort (AUC = 0.957). (**i**) Proportion of samples falling within the FGA groups for bivalency quartiles. Boxplots of the number of **(j)** SNVs, **(k)** tumor size and **(l)** mean expression of SS18::SSX target genes for the upper and lower bivalency quartiles. * indicates *P*-value < 0.05 for a Welch two-sample t test.

**Supplemental Figure 5.**
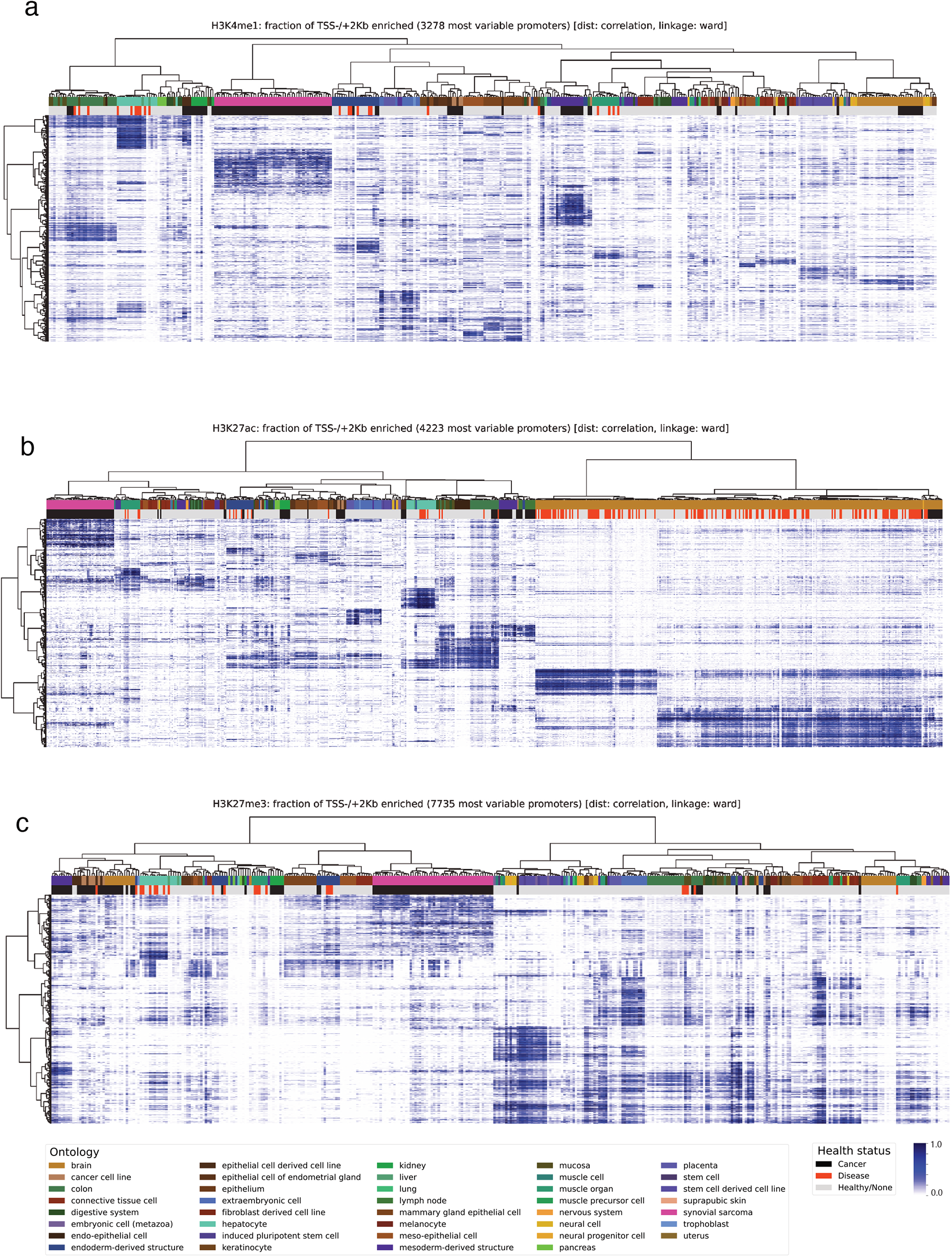
Synovial sarcoma has distinct promoter associated histone modification occupancy. Unsupervised hierarchal clustering (distance = 1 - spearman correlation, ward.D2 clustering method) of the proportion of promoter regions marked by (**a**) H3K4me1, (**b**) H3K27ac or (**c**) H3K27me3 in primary SyS (pink) and other normal and diseased tissue from the International Human Epigenome Consortium (IHEC).

**Supplemental Figure 6.**
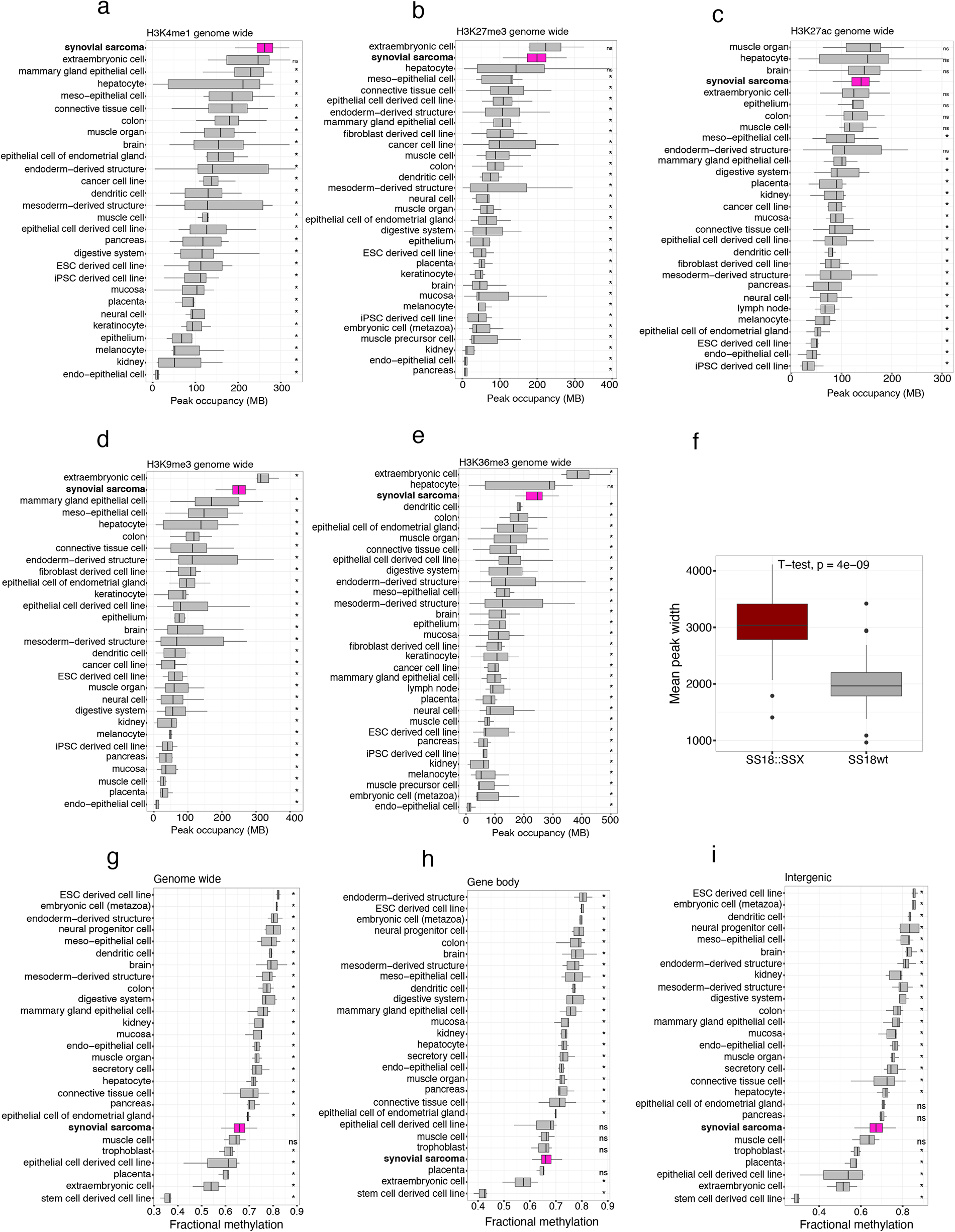
Synovial sarcoma has unique genome wide occupancy patterns of epigenetic marks. Genome wide occupancy of the histone marks **(a)** H3K4me1, **(b)** H3K27me3**, (c)** H3K27ac**, (d)** H3K9me3 and **(e)** H3K36me3 in SyS and control tissues. **(f)** H3K4me3 mean width of peaks overlapping SS18::SSX or SS18 wt binding sites. Mean fractional methylation of SyS and control tissues (**g**) genome wide, (**h**) in gene bodies and (**i**) intergenic regions. * indicates *P*-value < 0.05 using pairwise Wilcoxon signed-rank test.

**Supplemental Figure 7.**
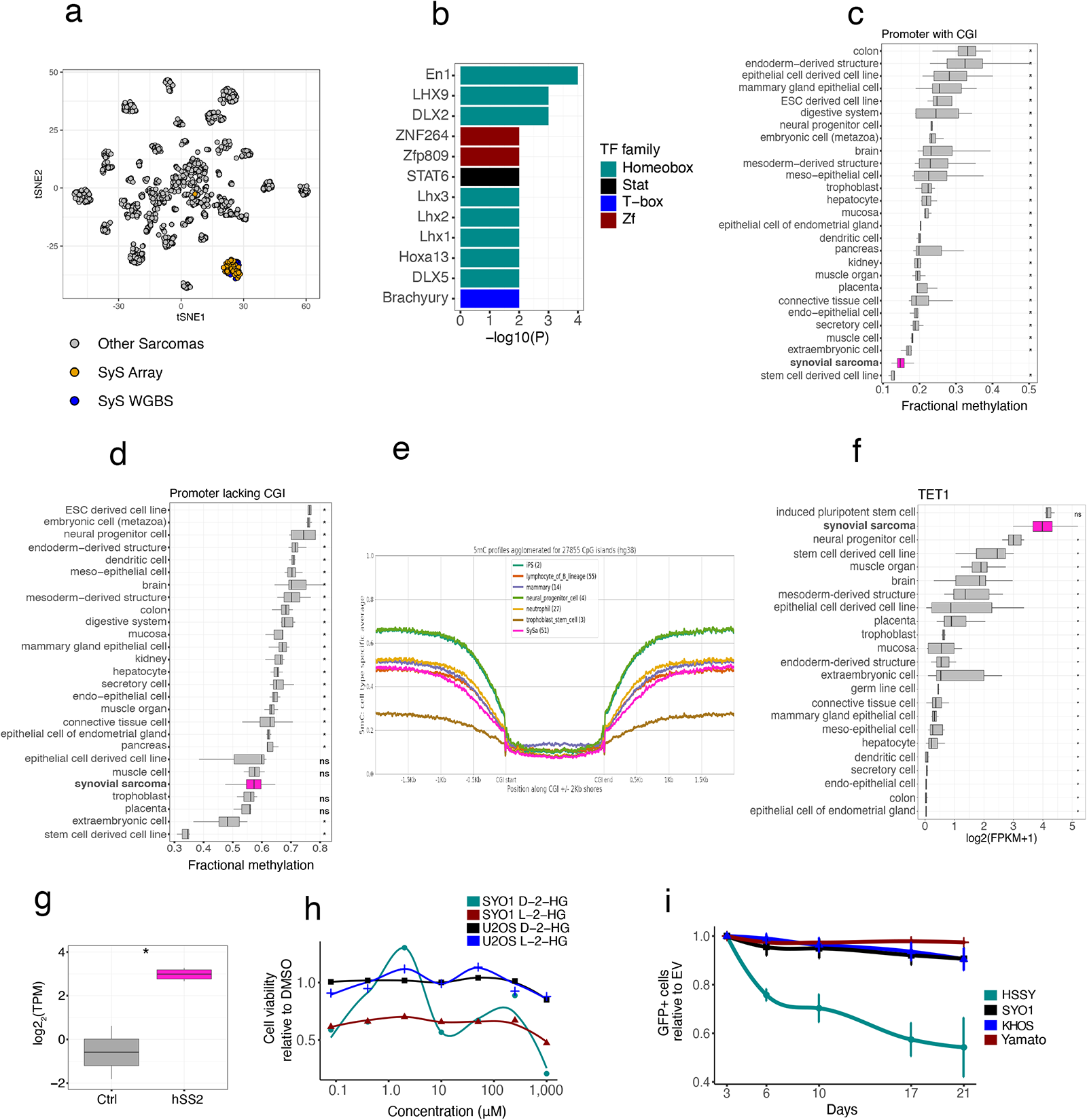
Synovial tumors have hypomethylated CGIs but are not sensitive to TET1 inhibition. **(a)** t-SNEA of beta values and fractional methylation values of SyS and other sarcoma subtypes. **(b)** Motif enrichment analysis in differentially hypomethylated regions. Mean methylation level of SyS and IHEC control tissues in **(c)** promoters harboring or (**d**) lacking CGIs. (**e**) Fraction methylation surrounding promoter bound CGIs in SyS and control tissues. (**f**) Expression of the *TET1* gene in SyS and IHEC control tissues. (**g**) Expression of the *TET1* gene in mouse and control samples. (**h**) Cell viability assay in SyS (HYSSII and SYO1) and Osteosarcoma (U2OS) cells upon 7-day treatment with 2-Hydroxyglutarate. (**i**) Cell competition assay performed in the synovial sarcoma lines HSSYII, SYO1, and Yamato transduced with an empty sgRNA as control or with guides targeting TET1. * indicates *P*-value < 0.05 using pairwise Wilcoxon signed-rank test.

**Supplemental Figure 8.**
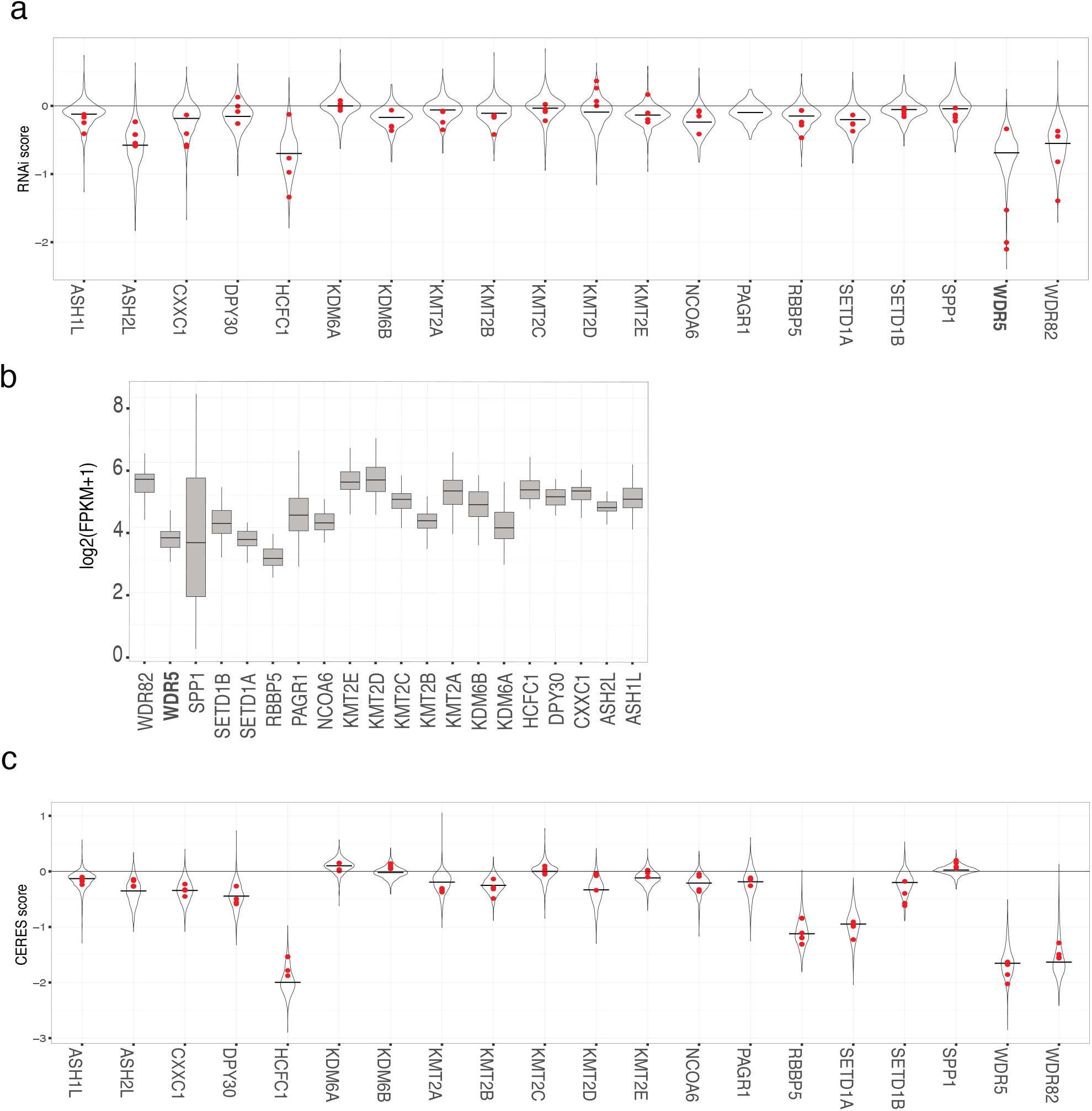
WDR5 is a selective vulnerability in synovial sarcoma. (**a**) Interrogation of RNAi viability scores for a panel of MLL and COMPASS complex members in SyS versus all other cell lines presented in the DepMap database identifies WDR5 as a selective vulnerability. Red dots indicate SyS cell lines and the black line within the violin plot represent the mean score for all cell lines. (**b**) Expression of the MLL and COMPASS complex members in SyS. (**c**) CRISPR CERES scores for a panel of MLL and COMPASS complex members in SyS versus all other cell lines presented in the DepMap database.

## Methods

### Biospecimens

82 fresh frozen primary SyS tumors were obtained from Vancouver General Hospital (Vancouver, BC), Mt. Sinai Hospital (Toronto, ON), Memorial Sloan Kettering Cancer Centre (New York City, NY) and Lund University Hospital (Lund, Sweden). Experiments involving patient tissues were carried out using protocols approved by the University of British Columbia Research Ethics Board (REB#: H18-00524, H18-02239, H18-02391, H12-01767). All cases had independent confirmation of *SS18::SSX* gene rearrangements by NanoString, RT–PCR or conventional cytogenetics during their clinical diagnostic work-up or central review. A single sarcoma containing a *BCOR::CCNB3* fusion with histologic overlap with SyS was included as a comparator. A detailed list of the tumors clinicopathological features are provided in **Table S1**.

### Total Nucleic Acid Extraction

Working on dry ice, two equal-size of approximately 3mm cube pieces were cut from each flash frozen primary SyS human tumor tissue without thawing and placed in separate 1.5mL LoBind tubes (Eppendorf, 022431021). Tissues were either kept on dry-ice or stored at -80°C for future processing. Bel-Art Liquid Nitrogen Cooled Mini Mortar and Pestle (VWR, 89233-994) were pre-chilled using LN2 (mortar) and dry ice (pestle). The tube containing frozen tissue was transferred from dry ice, placed inside the mortar, and cold ground using chilled pestle without direct contact with liquid nitrogen. Pulverized, still frozen tissue, was then quickly resuspended in the appropriate buffer (either ChIP-seq lysis buffer (see below) for the tissue entering NdChip-seq or Buffer RLT Plus (Qiagen, 80204) with B-mercaptoethanol (Sigma, M3148) for the tissue destined for dual RNA/gDNA extraction). Post pulverization, one tumor tissue piece was used for dual RNA/gDNA extraction using a combination of mirVana miRNA Isolation Kit (ThermoFisher, AM1560) and AllPrep DNA/RNA Mini Kit (Qiagen, 80204). Genomic DNA was eluted 6 times in ultrapure water (Thermofisher, 10977) for maximum recovery, concentrated using Vacufuge Centrifuge Concentrator (Eppendorf, 022820168), buffer adjusted to 10mM Tris-HCL, and stored at 4°C. Based on gDNA yield (Qubit dsDNA HS Assay, Thermofisher Q32854), the cellularity of the second matching tissue piece was estimated and used to determine the amount of chromatin entering NdChip-seq. Extracted gDNA was used for methylation profiling assays (PBAL, WGBS) and whole genome sequencing (WGS).

Total RNA was eluted 3 times in nuclease free water (Thermofisher, AM9937), ethanol precipitated, resuspended in 1/20 solution of SUPERaseIn RNase Inhibitor (Thermofisher, AM2696), and stored at -80°C. Total RNA was DNAse I treated (NEB, M0303), purified using in-house prepared magnetic bead solution (1M NaCL, 20% PEG, Sera-Mag Speedbead (Fisher Scientific, 09981123)), checked for quality and quantified using Agilent Bioanalyzer RNA Analysis (Agilent, 5067-1511). Extracted total RNA was used for ribodepleted, strand specific RNA-seq and Nanostring Assay.

### RNA Sequencing

RNA Sequencing (RNA-seq) was performed using a ribo-depleted, strand specific RNA-seq protocol as previously described^59^. 1uL of 1/100 dilution of ERCC RNA Spike-In Mix I (Thermofisher, 4456740) was added to all samples prior to library generation. Individually indexed libraries were pooled and sequenced on an Illumina platform (see **Supplemental Table 6**) to depth of 200M PE reads following the manufacture’s protocols (Illumina, Hayward CA.).

Paired-end reads were trimmed to 75-bp and processed through the grape-nf analysis pipeline (https://github.com/guigolab/grape-nf ) according to IHEC recommendations (https://github.com/guigolab/grape-nf/blob/master/ihec-setup.md). Chimeric transcripts were identified using STAR-Fusion (version 1.6.0)^60^ using default settings. Genes were annotated using Ensembl GRCh38 (hg38) version 94.

Differential gene expression analysis was performed using DEseq2^61^ (v1.26.0) using an FDR < 0.05 and FC > 2 as a cut-off. Single-sample Gene Set Enrichment Analysis (ssGSEA) was performed using GSVA^62^. The enrichment scores were calculated for SyS specific gene set defined by Jerby-Arnon *et al*^44^. Transcriptome deconvolution was performed with CIBERSORTx Fractions (https://cibersortx.stanford.edu/, absolute mode and 100 permutations) to estimate immune cell composition. Unsupervised hierarchical clustering was performed on the top 15 % most variably expressed genes based on FPKM values. Dendrogram was generated using pvclust (v2.2.0) (distance = 1 - spearman correlation, ward.D2 clustering method, bootstrap = 1000).

Isoform analysis was completed using the Mixture of Isoforms (MISO)^63^ (0.5.4). MISO settings were adjusted to analyze paired end reads (strand = fr-firststrand). Gene predictions for SS18::SSX were downloaded from the UCSC genome browser (https://genome.ucsc.edu/), and used to make the annotation file gff_make_annotation (--flanking-rule commonshortest --genome-label hg38). MISO was run (miso --run --read-len 75 --paired-end 190 70) and summarized (summarize_miso --summarize-samples) providing percentage spliced in values for the SS18::SSX isoforms. The reads specifically referring to exon 8 inclusion (chr18:26038555:26038659:-@chr18:26035831:26035923: @chr18:26035005:26035127:-) were compared between samples.

### Whole Genome Sequencing

Genomic DNA (gDNA) extracted from primary tumors (see above) and corresponding normal gDNA (blood or normal tissue) was subjected to PCR free whole-genome sequencing (WGS) as described^64^. Samples were uniquely indexed, pooled and sequenced on Illumina NovaSeq6000 as detailed in **Supplemental Table 6**. The resulted reads were aligned to hg38 using BWA and processed through the GATK v 4.2.0 pipeline and SNVs and INDELS were called using two different variant callers. Mutect (v2.2.1) SNV and INDEL predictions which passed all mutect filters (FilterMutectCalls) and were not found to be recurrent in a panel of normal were kept.

Strelka (v2.9.10) SNV and INDEL predictions which passed all strelka filters were also kept. The Mutect and Strelka calls retained after filtering were intersected and events called by both Mutect and Strelka were kept. Additionally, INDELS that were called by Strelka only but scored higher than a minimum threshold (QSI>=50) were also kept. The filtered VCF files were converted to MAF files using vcf2maf (v1.6.16) and variants were annotated using the Ensembl Variant Effect Predictor (VEP). CNV calling was performed on tumor bam files using CNVkit (v0.9.6) with the “–method wgs” option^65^. The copy-number reference was compiled using the corresponding normal samples. MAFtools^66^ (v2.10.5) was used for the visual representation of both missense mutations (oncoplot) and CNVs (heatmaps).

### Chromatin Immunoprecipitation Sequencing

Chromatin immunoprecipitation sequencing (ChIP-seq) was performed from frozen pulverized tumour tissue (as described above) using a nucleosome density protocol (NdChip-seq) as previously published^67^ for 8 histone modifications: H3K27ac (Hiroshi CMA309-IgG1), H3K27me3 (Diagenode C15410195), H3K4me3 (Cell Signaling #9751), H3K4me1 (Diagenode C15410037), H3K36me3 (Diagenode C15410192), H3K36me2 (abcam ab176921), H2AK119Ub (Cell Signaling Technology, 8240S) and H3K9me3 (Diagenode C15410056) using 100ng of MNase I digested chromatin per IP. Illumina sequencing libraries were generated by end repair, 3’ A-addition, and Illumina sequencing adaptor ligation (New England BioLabs, E6000B-10) as previously published^67^. Libraries were PCR amplified (8 cycles) using indexed primers, pooled, and sequenced on Illumina HiSeqX/ NovaSeq6000 , with read length PE75/PE150 to debth of 50M PE reads for narrow marks and 100M PE reads for broad marks and input (see **Supplemental Table 6**). SyS ChIP-seq libraries, as well as the data from normal and disease tissues obtained from the International Human Epigenome Consortium^29^ (IHEC), were uniformly processed according to the IHEC standardized workflow using the wrapper script (https://github.com/IHEC/integrative_analysis_chip). Read-depth normalized Bigwig files were generated using deepTools (3.3.0) bamCoverage (--normalizeUsing RPKM --ignoreDuplicates -- samFlagExclude 1028 --minMappingQuality 5 --binSize 20 --extendReads) (https://deeptools.readthedocs.io/en/develop/). Bigwigs were visualized using the UCSC genome browser (https://genome.ucsc.edu/). Read density was calculated using deepTools (3.3.0) multiBigwigSummary from normalized bigwig files. PyGenomeTracks (v3.7) was used to visualize MACS2 peak calls over specific genomic locations^68^.

Unsupervised clustering of SyS samples based on histone modifications was performed genome wide. Firstly, the genome (autosomes only) was binned into 500 bp bins and the proportion of each bin marked by a MACS2 peak for each histone modification was calculated. The most variable 60k bins across the tumors (approximately the top 1% most variable bins) were used for the unsupervised hierarchical clustering (distance = 1 - spearman correlation, ward.D2 clustering method, bootstrap = 1000). DiffBind (version 2.14.0) was then used to identify differentially bound regions between subgroups (FDR < 0.05, FC > 1). Distance from peak to closest TSS was calculated using “bedtools closest” and the empirical cumulative distribution was computed with stat_ecdf (https://github.com/tidyverse/ggplot2/blob/HEAD/R/stat-ecdf.R).

Bivalently marked regions were identified by intersecting MACS2 peak calls from H3K4me3 and H3K27me3 using bedtools (v2.30.0). Bivalently marked promoters were called by intersecting the bivalent regions with promoter regions (+- 2kb from TSS) of protein coding genes.

Unsupervised clustering of SyS and IHEC samples together was performed for promoter associated histone modifications. The a fraction of a promoters (TSS -2/+0.3Kb for H3K4me3 and -2/ +2Kb for H3K4me1, H3K27ac, and H3K27me3) overlapping MACS2 enriched regions was calculated for all protein coding genes (Ensembl v94) using bedtools (v2.30.1). Genes with low signal (<0.01) for all data sets were excluded and only genes with high variability between samples (std > 1.25 mean) were considered. After this selection, depending on the histone modification we ended up with the ∼10-15% most variable promoters. We used Python scipy package for the clustering (and seaborn clustermap for visualization). Hierarchical clustering of both rows and columns was performed with a Pearson correlation as a similarity measure and Ward linkage was applied.

### Public ChIP-seq data

Raw sequencing files (fastq files) corresponding to ChIP-seq from human cell lines (Aska and SYO1)^25^ and mouse cell line C3H10T1/2^24^ were downloaded from Gene Expression Omnibus (GEO) accession number GSE108028 and GSE 148724, respectively. Reads were aligned to hg38/mm10 using BWA-MEM (0.7.6a)^69^. BAM files were sorted using Sambamba (0.5.5)^70^ and duplicates were marked using Picard Tools (1.52) MarkDuplicates (http://broadinstitute.github.io/picard/). Peaks were called using MACS2 (2.1.1.20160309) (FDR < 0.05)^71^. SS18::SSX target sites were determined by overlapping SS18 peak calls in the shCt condition from both Aska and SYO1 and then removing SS18 regions found in the shSSX conditions.

### Whole Genome Bisulfite Sequencing

Whole genome DNA methylation data was generated using two methods: Post-Bisulfite Adapter Ligation (PBAL) and Whole Genome Bisulfite Sequencing (WGBS). To track the efficiency of bisulfite conversion, 1% of an equal molar mix of unmethylated lambda DNA (Promega, D1521) and fully methylated T7 was spiked into genomic DNA. PBAL libraries were generated as previously described^72^ with the following modifications: starting material was 100ng of gDNA, one round of DNA generation/random priming was used post sodium bisulfite conversion, PCR was limited to four rounds, and after PCR a size selection step was used to enrich for larger fragments of DNA. Uniquely indexed Illumina libraries generated by both methods were pooled and submitted for Illumina sequencing following the manufacture’s protocols (Illumina, Hayward CA.).

PBAL and WGBS reads were aligned to hg38 and processed using the gemBS pipeline (version 3.5.0)^73^. Methylation values were called for each CpG. To calculate fractional methylation for regions of interest, a weighted average was used based on the methylation value and coverage of CpGs. A minimum coverage of 3 was used to filter regions. CNV calling was performed on tumor bam files generated by gemBS using CNVkit (v0.9.6) with the “–method wgs” option^65^.

### DNA methylation array analysis

Processed DNA methylation array data (beta values) for 1505 sarcoma samples^50^ was downloaded from Gene Expression Omnibus using accession number GSE140686. Differentially methylated probes (DMPs) were called using ChAMP (version 2.20.1) and the ProbeLasso algorithm^74,75^ was used to identify differentially methylated regions (DMRs). Unsupervised clustering was performed on the combined public and in-house generated sarcoma datasets. The WGBS data was filtered to only contain CpG calls from Illumina array probe positions and then merged with public array dataset. We performed unsupervised non-linear dimension reduction according to the original publication, first the 10,000 most variable probes according to standard deviation were selected and the t-SNE plot was then computed with Rtsne (v0.16) (3000 iterations and a perplexity value of 30).

### Survival analysis

Metastasis-free survival (MFS) was calculated from the date of diagnosis until the date of metastatic occurrence or death from any cause and follow-up was measured from the date of diagnosis to the date of latest follow-up for event-free patients. Survival was calculated using the Kaplan–Meier method and curves were compared with the log-rank test using the survminer package (v0.4.9).

### NanoString

A custom CodeSet to target 50 genes of interest was designed and provided by NanoString technologies for the nCounter system (**Supplemental table 5**). RNA extraction, sample preparation and probe–RNA complex quantification was performed according to manufacturer’s recommendations as previously described^45^. NanoString nSolver software’s default postprocessing settings were applied to generate normalized gene counts.

A random forest model was trained on a set of 31 samples using the R statistical software package randomForest^76^. The model was tuned using tuneRF and evaluated using receiver operating characteristic (ROC) curves and Area Under the Curve (AUC) metrics using the ROCR package (v1.0.11). The model was then applied to predict the bivalency group of the remaining 51 samples for validation.

### Genomic region annotation

Gene ontology analysis was performed using Metascape^77^. Enriched terms are ranked by -log(*p*-value). Enriched transcription factor binding motifs were identified using HOMER^78^ (version 4.9) findMotifsGenome.pl.

### Western Blots

Two different synovial sarcoma cell lines were used; HSSYII (RIKEN, Saitama, Japan), and SYO-1 (Dr. Akira Kawai, National Cancer Centre Hospital, Tokyo, Japan). An osteosarcoma cell line, U2OS, was used as a non-synovial sarcoma comparator. All cell lines were cultured at 37°C with 5% CO_2_ in Gibco RPMI 1640 Media (cat. 11875093) supplemented with 10% fetal bovine serum (sigma). Cells were cultured in appropriate conditions for 72 hours being washed, pelleted and lysed using RIPA Lysis buffer (Milipore cat. 20-188) with a protease inhibitor cocktail (Sigma cat. 539134). Cells were centrifuged and the supernatant was collected, and protein amounts were quantified using the Pierce BCA Protein Assay Kit (cat. 23225). Protein amounts were normalized to ensure equal protein loading into each lane. Lysates were then prepared with NuPAGE 4X LDS Sample Buffer (cat. NP0008), and NuPAGE 10X Sample Reducing Agent (cat. NP0004). Samples were heated to 70°C for 10 minutes and loaded into NuPAGE 4-12% Bis-Tris Gels. The Thermofisher MES nuPAGE system was used for running and transferring the western, as per the manufactures instructions. Gels were transferred onto Biorad Nitrocellulous Membrane (cat. 1620112). Membranes were blocked in the LiCor Intercept blocking buffer (cat. 927-70001). Antibodies used include Cell Signalling SS18::SSX (cat. 72364), Santa Cruz GAPDH (cat. sc-47724), IRDye800CW (cat. 925-32210), IRDyeG80RD (cat. 925-68071). Blots were imaged using the LiCor Oddessy Imager and intensity was quantified with ImageJ.

### Tissue microarray immunohistochemical imaging

Tissue microarray (TMA) construction from anonymized patient primary surgical excision specimens was performed under protocols H18-00524 and H18-02391, approved by the Clinical Research Ethics Board of the University of British Columbia and BC Cancer. AIRID1A, BRG1, PBRM1, BRD9 and SS18::SSX immunohistochemistry was performed on a 4-µm section of a formalin-fixed, paraffin-embedded (FFPE) human TMA consisting of 37 synovial samples from Vancouver General Hospital. Cases were included as 0.6 mm patient sample cores in duplicate. Additionally, 16 individual whole tissue sections were cut from paraffin and mounted on slides to create 4-µm sections that were stained individually. The assays were run with the following conditions via a Leica BOND RX (Leica Biosystems). Heat-induced epitope retrieval was performed using citrate-based BOND Epitope Retrieval Solution 1 (Leica Biosystems) for BRG1, PBRM1, BRD9, and S18::SSX, for 10min, 20min, 10 min, and 20min, respectively. The EDTA-based BOND epitope Retrieval 2 was used for ARID1A for 10 min. The primary antibodies ARID1A (Abcam, ab182560), BRG1 (Abcam, ab110641), BRD9 (Thermo Fisher Scientific, 24785-1-Ap), SSX::SS18 (Cell Signaling Technology, 72364S), PBRM1 (Cell Signaling Technology, 81832) were incubated at ambient temperature at 1:3000 for 30 min,1:2000 for 30 min, 1:200 for 30min, 1:300 for 15min, and 1:100 for 30min, respectively. Staining was visualized using the BOND Polymer Refine Detection kit (Leica Biosystems, DS9800), which includes a 3,3′-diaminobenzidine (DAB) chromogen and hematoxylin counterstain. TMA virtual slide scans were then generated on a Leica Aperio AT2 (Leica Biosystems) at ×40 magnification. Each individual patient sample core was analyzed using HALO and HALO AI (Indica Labs), which required user annotated training data to develop an artificial intelligence segmentation network for nuclear identification. The TMA module was implemented to extract individual patient core images from the TMA whole slide scan. The Multiplex IHC module was trained to identify DAB staining using representative pixels for delineation from hematoxylin in order to determine average DAB nuclear optical density.

### RNA Interference

Cells were cultured in Gibco RPMI 1640 Media (cat. 11875093) and plated in 6-well plates at ∼60%. The following day, cells were transfected with 9uL of RNAiMAX lipofectamine (cat. 13778150), 10pmol of pooled siRNA, and 300uL of opti-MEM (cat. 13985062), added dropwise to the wells. Cells were passaged and re-transfected every 3 days. FAM tags were included on siRNA constructs to confirm transfection efficiency. Western blots were run to confirm knockdown and cell viability and counts were recorded. Duplex oligo constructs were ordered from Integrated DNA Technologies (IDT) and had the following sequences: siSSX2 sense: /56-FAM/ rCrArA rGrArA rGrCrC rArGrC rArGrA rGrGrA rATT, antisense: rUrUrC rCrUrC rUrGrC rUrGrG rCrUrU rCrUrU rGTT as previously described (Laporte *et al*., 2017; Lubieniecka *et al*., 2008). The Invitrogen Silencer™ FAM-labeled Negative Control No. 1 siRNA (cat. AM4620) was used as a control in all experiments.

### Cell viability assay

Cells were seeded in 96-well plates (HSSYII at 1 × 10^3^ cells/well, SYO-1 at 1 × 10^3^ cells/well and U2OS at 5 × 10^2^ cells/well) and treated in triplicate at indicated doses (range 0.016 – 50 uM) of the WDR5 inhibitor, OICR-9429. The drug and media were refreshed on day 3 and cell viability was assessed in the cell lines as compared with the vehicle condition (0.1% DMSO) at 7 days post treatment using MTS reagent (Promega, Madison, WI, USA). IC_50_ values were determined in the cell lines by dose-response curves calculated using the drc (v3.0.1) R package.

### Cell competition assays

HSSYII, SYO1, Yamato and KHOS Cas9 cells were transduced with an empty plasmid (empty vector) or a plasmid containing sgRNA targeting WDR5 or TET1. Infections were done with a virus dilution of 1:10 to obtain an infection efficiency of around 70–80%. Infected cells become GFP+ due to the backbone of the sgRNA. The cells were then cultured over a period of 25 days, and the percentage of GFP+ cells was measured using a Fortessa fluorescence-activated cell sorting (FACS) machine.

